# Impaired DNA damage response and inflammatory signalling underpins hematopoietic stem cell defects in *Gata2* haploinsufficiency

**DOI:** 10.1101/2024.08.20.608056

**Authors:** Ali Abdelfattah, Ahmad Habib, Leigh-anne Thomas, Juan Bautista Menendez-Gonzalez, Alhomidi Almotiri, Hind Alqahtani, Hannah Lawson, Sarab Taha, Millie Steadman, Radhika Athalye, Alex Gibbs, Hamed Alzahrani, Alice Cato, Peter Giles, Alex Tonks, Ashleigh S. Boyd, Kamil R. Kranc, Neil P. Rodrigues

**Author notes:** Corresponding author: Neil P. Rodrigues, DPhil European Cancer Stem Cell Research Institute Cardiff University, School of Biosciences, Hadyn Ellis Building Maindy Road Cardiff, CF24 4HQ UK.

## Abstract

Clinical *GATA2* deficiency syndromes arise from germline haploinsufficiency inducing mutations in *GATA2*, resulting in immunodeficiency that evolves to myelodysplastic syndrome (MDS)/acute myeloid leukemia (AML). How *GATA2* haploinsufficiency disrupts the function and transcriptional network of hematopoietic stem/progenitors (HSCs/HSPCs) to facilitate the shift from immunodeficiency to pre-leukemia is poorly characterised. Using a conditional mouse model harboring a single allele deletion of *Gata2* from the start of HSC development *in utero*, we identified pervasive defects in HSPC differentiation from young adult *Gata2* haploinsufficient mice during B-cell development, early erythroid specification, megakaryocyte maturation to platelets and inflammatory cell generation. *Gata2* haploinsufficiency abolished HSC self-renewal and multi-lineage differentiation capacity. These functional alterations closely associated with deregulated DNA damage responses and inflammatory signalling conveyed from *Gata2* haploinsufficient HSCs. We identified genetic interplay between *Gata2* and *Asxl1*, a driver of DNA damage and inflammation and, notably, a recurrent secondary mutation found in *GATA2* haploinsufficiency disease progression to MDS/AML. shRNA mediated knockdown of *Asxl1* in *Gata2* haploinsufficient HSPCs led to an enhanced differentiation block *in vitro*. By analysis of HSCs from young adult compound *Gata2/Asxl1* haploinsufficient mice, we discovered hyperproliferation of double haploinsufficient HSCs, which were also functionally compromised in transplantation compared to their single *Gata2 or Asxl1* haploinsufficient counterparts. Through both *Gata2/Asxl1* dependent and unique transcriptional programs, HSCs from compound *Gata2/Asxl1* haploinsufficient fortified deregulated DNA damage responses and inflammatory signalling initiated in *Gata2* haploinsufficient HSCs and established a broad pre-leukemic program. Our data reveal how *Gata2* haploinsufficiency initially drives deregulation of HSC genome integrity and suggest the mechanisms of how secondary mutations like *ASXL1* take advantage of HSC genomic instability to nurture a pre-leukemic state in *GATA2* haploinsufficiency syndromes.

## Introduction

Blood and immune cells with wide-ranging purpose including immune defence, blood clotting and oxygen supply to tissues, originate from scarce, self-renewing hematopoietic stem cells (HSCs) housed in the bone marrow (BM) (1). Cell intrinsic genetic and epigenetic mechanisms combine with cell extrinsic signalling relayed from the BM niche to regulate HSC cell cycle and apoptotic fates that assure maintenance of genomic stability in HSCs for life (1, 2). However, mechanisms critical for maintaining HSC integrity - governed principally by nuclear transcription factors (TFs) – can become corrupted through the acquisition and functional propagation of genetic and epigenetic mutations in HSCs and their descendants, progenitor cells (HSPCs), increasing the probability of immunodeficiency, BM failure syndromes, hematologic malignancies, and other non-hematopoietic co-morbidities, such as cardiovascular disease (1, 2). Aging is a major determinant for the emergence of these conditions, including leukemia. Yet leukemia can also arise in children and at a relatively young age through inherited genetic predisposition, exemplified by hereditary myelodysplastic syndrome (MDS)/acute myeloid leukemia (AML) caused by mutations in *RUNX1*, *CEBPA* or *GATA2* TFs (3).

As a member of the GATA (Guanine/Adenine/Thymine/Adenine) family of zinc-finger (ZnF) TFs that regulate gene expression by binding to the DNA sequence WGATAR (W:A/T, R:A/G) (4, 5), *Gata2* is most prominently expressed in the HSC compartment and, reflecting this expression pattern, extensive experimental modelling using loss and gain of function approaches has identified *Gata2* as an indispensable regulator of HSC development from hemogenic endothelium and of adult HSC maintenance in the BM (6–10). Though expression of *Gata2* variably diminishes during differentiation to select hematopoietic lineages, nonetheless it has defined roles in lineage specification, including granulocyte-macrophage progenitor generation and terminal differentiation of megakaryocytes, basophils, and mast cells (6, 11–14). These data highlight *Gata2* as a crucial regulator of HSC generation, adult HSC maintenance, and differentiation to select hematopoietic lineages.

Mirroring the similar functional outcomes of down-regulating and up-regulating GATA2 expression level in HSPCs in loss and gain of function studies respectively (9, 11, 15, 16), deregulation of *GATA2* expression in the clinical setting has been shown to have both tumour suppressor and oncogenic roles in AML. Overexpression of *GATA2* confers poor prognosis in 25% of all AML cases, implying an oncogenic role for GATA2 in AML (17). Studies where *Gata2* was genetically deleted in mouse models of AML lead to enhanced apoptosis specifically in the therapy resistant leukemic stem cell (LSC) compartment and reactivation of myeloid differentiation, suggesting that the *Gata2* transcriptional network may be an exploitable therapeutic vunerability in AML (8). As proof of concept supporting this notion, pharmacologic inhibition of GATA2 in combination with AML chemotherapy has been shown to enhance apoptosis of AML cells (18, 19).

In striking contrast, a tumour suppressor role for *GATA2* has also been established in MDS/AML. Hereditary *GATA2* haploinsufficiency mutations are observed throughout the *GATA2* locus, yet approximately 70% of mutations arise in the DNA binding zinc finger domains (20, 21), disrupting GATA2 chromatin binding activity and the expression of normal GATA2 target genes in hematopoiesis (22, 23). Germline *GATA2* haploinsufficiency mutations are largely categorised into four groups: (i) truncated mutations (i.e. nonsense, in-frame-deletion and in-frame-insertion mutations), which represent approximately 60% of *GATA2* haploinsufficiency cases; (ii) missense mutations reported in approximately 30% patients; (iii) non-coding mutations in intron-4 (+9.5kb cis-element), which represent about 10% of cases; and (iv) sporadic cases of whole-locus deletions, N/C terminals, and untranslated region (UTR) regions (20, 21, 24). These loss of function *GATA2* mutations, which largely result in loss of function, initially lead to primary immunodeficiency disorders such as MonoMAC and dendritic cell, monocyte, B/NK lymphoid (DCML) syndromes, which are characterised by a severe reduction in peripheral monocytes, CD4^+^ lymphocytes, B-cells, NK-cells, and dendritic cells, and Emberger syndrome, an autosomal dominant disorder characterised by primary lymphedema of the lower limbs, cutaneous-warts, and deafness (25–29). The development of Emberger syndrome in the context of *GATA2* haploinsufficiency reflects the role of *Gata2* in lymphatic vessel development during embryogenesis (28, 30, 31). *GATA2* haploinsufficiency driven MonoMAC, DCML and Emberger syndromes ultimately evolve to MDS/AML on acquisition of additional co-operating mutations, such as *ASXL1*, with variable latency and presentation (28, 32–34).

There are compelling imperatives to understand the mechanisms underscoring *GATA2* haploinsufficiency in HSCs/HPCs. First, widespread myeloid and lymphoid differentiation defects are observed in the setting of *GATA2*-haploisufficiency mediated MonoMAC/DCML immunodeficiency disorder and pan-cytopenia is similarly observed in Emberger syndrome (26, 27, 29, 32), suggesting a functional defect originating in the HSC/HSPC compartment; such postulate is supported by extensive modelling of haploinsufficiency in genetic mouse models, zebrafish models and shRNA mediated knockdown in cord blood HSPCs (8, 9, 11, 25, 35, 36). Furthermore, while most cases of MonoMAC evolve to myeloid neoplasms, some MonoMAC patients with *GATA2* deficiency advance to B-ALL (37–39). Second, variable patient disease presentation and latency, even for similar types of *GATA2* haploinsufficiency mutations, may be ascribed to the differential genetic/epigenetic impact of similar *GATA2* mutations occurring within the heterogeneous HSC/HSPC compartment (20, 21). Alternatively, variations in patient progression could be due the influence of secondary mutations that are adopted in HSCs/HSPCs from *GATA2* haploinsufficiency patients. Finally, HSC/HSPC derived pre-LSCs and LSCs in MDS/AML provide a reservoir of leukemic cells that are not cleared by current AML treatment regimens, forming the cellular basis for therapy resistance, relapse, and fatality in AML (33, 34). How the pre-LSC/LSC transcriptional programme mediated by *GATA2* haploinsufficiency shapes initiation, propagation, or maintenance of leukemia in these patients remains unclear. Understanding the functional and transcriptional network operating in *GATA2* haploinsufficient HSCs and the impact of secondary co-operating mutations in HSCs/HSPCs in *GATA2* deficiency is therefore required to conceive and identify targeted, non-toxic curative treatments rather than the prevailing, imperfect standard of care for these patients, allogeneic bone marrow transplantation (33, 34).

Germline and conditional mouse genetic models of *Gata2* haploinsufficiency have recapitulated the hereditary nature and select biological functions underlying clinical *GATA2* mutation driven syndromes that, in the vast majority of patients, correlates with *GATA2* haploinsufficiency and reduced *GATA2* expression (7, 9, 11, 36, 40). By breeding *Gata2* “floxed” allele mice *(Gata2^fl/fl^)* with constitutively active, hematopoietic-specific *Vav-Cre* mice (*Vav-iCre^+^*), we generated *Gata2^+/fl^; Vav-iCre^+^* mice which harbored a genetic deletion in a single *Gata2* allele from the onset of HSC emergence *in utero* at embryonic day 11. Through analysis of young adult *Gata2^+/fl^; Vav-iCre^+^* mice, in this report we sought to understand the hitherto poorly understood mechanistic impact of *Gata2* haploinsufficiency in HSCs/HPCs.

## Experimental Materials and Methods

### Animals

Generation of *Gata2^fl^*^/fl^, *Asxl1 ^fl/fl^*, and *Vav-iCre* mice has been previously described (41–43). *Gata2^fl^*^/fl^ mice were kindly donated by Dr Julian Downward, The Crick Institute, London. *Vav-iCre* mice were kindly gifted by Professor Kamil Kranc, Institute for Cancer Institute, London. *Asxl1 ^fl^*^/fl^, C57Bl/6J (CD45.2), and C57Bl/6J-SJL (CD45.1) mice were purchased from Jackson Laboratory. *Gata2^fl/fl^; Vav-iCre^-^* males were bred with *Gata2^+/+^; Vav-iCre^+^*females to obtain *Gata2^+/fl^; Vav-iCre^+^* (*Gata2* heterozygote) and *Gata2^+/fl^; Vav-iCre^-^* (control) mice. For double heterozygote mice of *Gata2* and *Asxl1*, *Gata2^+/+^; Vav-iCre^+^* females were intercrossed with *Gata2^+/fl^; Asxl1^+/fl^, Vav-iCre^-^* males to generate an experimental cohort of *Gata2^+/fl^; Vav-iCre^+^* (single *Gata2* heterozygote), *Asxl1^+/fl^; Vav-iCre^+^* (single *Asxl1* heterozygote), *Gata2^+/fl^, Asxl1^+/fl^, Vav-iCre^+^* (double heterozygote mice), and *Gata2^+/fl^, Asxl1^+/fl^, Vav-iCre^-^*(control). All experimental mice were age and sex matched in a C57BL/6 genetic background and were assessed at 8-12 weeks old. Mice were bred and housed at Heath Park Unit, Cardiff University. All animal procedures were carried out in accordance with the Animals (Scientific Procedures) Act 1986, the United Kingdom Home Office authorization.

### Peripheral blood analysis

Peripheral blood (PB) was collected by the lateral tail venesection using EDTA microvette capillary collection tubes (Sarstedt). The HemaVet (Drew Scientific) machine was used to automatically count hematological indices according to standard manufacturer’s instruction. For immunophenotypic analysis of PB, red blood cells (RBCs) were lysed using ammonium chloride lysis buffer (STEMCELL Technologies).

### Preparation of bone marrow (BM), spleen, and thymus cells

Two femurs and tibias bones were dissected from each mouse removing muscles and skin tissues and crushed in phosphate-buffered saline (PBS) with 2% fetal bovine serum (FBS) (PBS 2%FBS) (Gibco) using a pestle and mortar, and cell suspension collected through a 70 μm cell strainer to remove small bone fragments (Miltenyi Biotec). Spleen and thymus tissues were gently minced in PBS with 2% FBS solution through a 70 μm cell strainer using a 2 mL syringe-plunger. Harvested cells from BM, spleen, and thymus were counted utilizing the BD Accuri^TM^ machine (BD Biosciences).

### Flow cytometry

Antibody staining was carried out according to the manufacturer’s instruction. For immunophenotypic analysis of lineage positive cells, 2×10^5^ cells of PB, BM, spleen, and thymus were harvested as described above and stained with appropriate antibody cocktails targeting myeloid cells (Mac1, Gr1), erythrocytes (Ter119, CD71), mature megakaryocytes (CD41, CD42d), B-cells (B220), T-cells (CD3, CD4, CD8),inflammatory monocytes (Mac1, Ly6C, Ly6G, CD115), and B lymphocyte precursors (CD19, CD43, IgM, BP1) (BioLegend) and incubated for 30 minutes at 4°C. Stained cells were then washed with PBS 2%FBS solution, resuspended in 200 μL of PBS 2%FBS with 2 μL of 20 μg/mL 4, 6-Diamidino-2-Phenylindole, Dihydrochloride (DAPI, Molecular Probes) with a final concentration of 0.2 μg/mL to determine live cells and subsequently analyzed by flow cytometry. For immunophenotypic analysis of BM HSPCs and committed progenitors, 10×10^6^ cells were stained in PBS with 2% FBS at 4°C for 30 minutes with following antibodies: CD16/CD32 (Fc Block, only for HSPCs), a biotinylated lineage-positive cocktail (CD3, CD4, CD8a, B220, Mac1, Gr1, Ter119), c-kit, Sca1, CD150, CD48, CD34, CD135, CD16/32, and CD127 (BioLegend, eBioscience). Stained cells were then washed and incubated with a streptavidin (BioLegend, eBioscience) conjugate for 20 minutes at 4°C. Next, cells were washed and subsequently resuspended in 600 μL of PBS 2% FBS with 6 μL of 20 μg/mL of DAPI (final concentration of 0.2 μg/mL). Lastly, the cells were filtered through a 30 μm Nylon filter (Sysmex) before being analysed. For FACS analysis of transplantation experiments, CD45.1 and CD45.2 antibodies were added to the staining mix of lineage positive cells, HSPCs, and committed progenitors. For apoptosis assay, BM HSPCs were stained with cell-surface fluorochrome cocktails as described above and then stained with conjugated annexin V (BioLegend) in 100 μL of annexin binding buffer (BioLegend) and incubated for 30 minutes at RT in the dark. Cells were next resuspended in 500 μL of annexin binding buffer with 6 μL of 20 μg/mL DAPI (final concentration of 0.2 μg/mL) and filtered through a 30 μm Nylon filter before flow-cytometric analysis. For Intracellular staining, after the extracellular staining as described above, cells were washed with cold PBS and fixed in 1% paraformaldehyde (ThermoFisher) for 20 minutes on ice and then permeabilized in 0.1% Saponin (Sigma) for 15 minutes on ice and intracellularly stained with conjugated Ki-67, γH2AX or isotype-control fluorochromes (BioLegend, Miltenyi Biotec, Cell Signaling Technology) and incubated for 30 minutes at 4°C and followed by flow cytometry. For Ki-67 assay, cells were incubated with 5 μg/mL DAPI for 5 minutes before being analysed. Flow cytometry was performed on LSR Fortessa^TM^ (BD, Biosciences) flow cytometer. Analysis of flow cytometric was performed using FlowJo 10.6.1 software (Tree Star). Flow cytometry data were analyzed as described previously (44–49).

### Magnetic c-kit^+^ cells enrichment and cell sorting

BM cells were enriched for c-kit positive cells using MACS (Miltenyi Biotec) according to the manufacturer’s protocol. BM cells were stained with anti-c-KIT magnetic beads (Miltenyi Biotec) and incubated for 20 min at 4°C in the dark on the rotating shaker. Enriched c-Kit^+^ cells were then stained as described above and utilized for LSK (Lin^−^Sca1^+^c-Kit^+^) or HSCs (LSK_CD150^+^CD48^-^) sorting. Stained Cells were isolated by FACS using a FACSAria^TM^ cell sorter (BD Biosciences).

### Transplantation experiments

CD45.2 BM cells were freshly harvested, enriched for a c-kit^+^ marker, stained with surface HSPC fluorochromes, and sorted for HSCs (Lin^-^c-kit^+^Sca1^+^CD48^-^CD150^+^) on FACS Aria as described above. CD45.1 mice (C57BL/6 SJL) were used as recipient hosts for all transplantation experiments. A day before transplantation, recipient hosts were lethally irradiated with two doses of 450 cGy divided by a 4-hour interval. For primary transplantation, 150 CD45.2^+^ HSCs from *Gata2^+/fl^; Vav-iCre^+^*, *Gata2^+/fl^; Vav-iCre^-^, Asxl1^+/fl^; Vav-iCre^+^*, *Gata2^+/fl^, Asxl1^+/fl^, Vav-iCre^+^,* or *Gata2^+/fl^, Asxl1^+/fl^, Vav-iCre^-^* mice were mixed with 2×10^5^ CD45.1^+^ competitor BM cells and transplanted via tail vein injection into lethally irradiated CD45.1^+^ host mice as described previously (50). For secondary transplantation, 200 CD45.2 HSCs were purified from *Gata2^+/fl^; Vav-iCre^+^* and control primary recipient mice at week sixteen after transplantation together with 2×10^5^ CD45.1 support BM cells and transplanted intravenously into secondary lethally irradiated CD45.1^+^ recipient hosts. Tail-vein bleeding was performed at different time points after transplantation to assess donor derived cells. Mice were sacrificed at week 16 post transplantation, and BM, spleen and thymus were harvested for immunophenotypic analysis.

### *In vitro* colony-forming cell assay

To quantify myeloid progenitor colonies, a total of 2×10^4^ live BM cells were plated in cytokines supplemented MethoCult M3434 media (STEMCELL Technologies) containing SCF, IL-3, IL-6, erythropoietin, insulin, and transferrin and incubated at 37°C and 5% CO_2_ as per the manufacturer’s instruction. Colonies were scored at day 10-12 post plating with a light microscope.

### Lentiviral production and transduction

Scramble (VB150923-10051), *mAsxl1-* sh1 (VB180515-1197ubr) and *mAsxl1-*sh2 (VB180515-1198gbn) vectors were purchased from VectorBuilder company. VSV-G and psPAX2 plasmids were kindly provided by Dr Kamil Kranc, Institute of Cancer Research, London. All DNA plasmids were purified using the EndoFree Plasmid Maxi-Kit (Qiagen) as per the manufacturer’s instruction. Lentiviral production was performed using calcium phosphate transfection method (51). Briefly, lentiviral vectors (scramble or *mAsxl1-* shRNA), psPAX-2 packaging plasmid, and VSV-G envelope plasmid were mixed with calcium chloride (Sigma) and 2x-HEPES buffered saline (Sigma) and incubated for 12-15 min at RT, and approximately 60-70% confluent human embryonic kidney 293T (HEK 293T, Cell Biolabs Inc) cells were transfected with lentiviral vectors. Lentiviral supernatant was collected 48 and 72 hours post transfection, filtered through a 0.45 μm filter (Sigma) and subsequently concentrated by ultracentrifugation using Optima XPN-80 Ultracentrifuge (Beckman Coulter) at 23,000 xg for 2 hours at 4° C. For LSK (Lin^−^ Sca-1^+^ c-Kit^+^) transduction, 5 ×10^4^ murine LSK cells were isolated as described above, pre-stimulated in IMDM Medium (StemCell^TM^ Technologies) supplemented with 100 ng/mL m-SCF (Peprotech), 100 ng/mL m-TPO (Peprotech) and 100 ng/mL mFlt3L (Peprotech) for 3-4 hours, transferred to pre-coated-retronectin (TaKaRA Bio) 96-well plate according to the manufacturer’s instruction with a multiplicity of infection (MOI) of 50 and incubated overnight at 37°C, 5% CO_2_. The next day, cells were washed in IMDM and plated with new IMDM supplemented with cytokines as described above and incubated at 37°C, 5% CO_2_ for 72 hours before analysing mCherry positive cells.

### Genomic PCR and reverse transcription (RT)-qPCR

Mouse genomic DNA was extracted from either mouse ear tissues (25 mg) or BM (up to 1×10^7^ cells) according to the manual of Isolate 1 Genomic DNA kit (Bioline). PCR Mix was prepared in a 25 μL total volume including; 12.5 μL of MangoMix (Bioline), 8.3 μL of nuclease free-water (Molecular Probes), and 0.1 μL of each forward and reverse primer (Sigma-Aldrich) and performed on 100^TM^ Thermal Cycler (Bio-Rad) using the following parameters: 94°C for 3 minutes; 35 cycles of 94°C for 1 minute, 60°C for 1 minute, 72°C for 1 minute; and 72°C for 10 minutes. *Gata2*-forward (5’ GCCTGCGTCCTCCAACA CCTCTAA 3’) and *Gata2*-reverse (5’ TCCGTGGGACC TGTTTCCTTAC 3’) primers. *Asxl1*-forward (5’ ACACCAACCAGCCGTTTTAC 3’) and *Asxl1*-reverse (5’ TCCTTGGATTTTTCTCAGCA 3’). Excised *Asxl1* allele was determined using *Asxl1*-forward (5’ ACGCCGGCTTAAGTGTACACG 3’) and *Asxl1*-reverse (5’ GACTAAGTTGCCGTGGGTGCT 3’). A 2% agarose gel with a safeview nucleic acid stain (NBS Biologicals) was used to detect DNA bands that were subsequently visualized by the ChemiDoc™MP imaging system (Bio-Rad). For RT-qPCR, total RNA was extracted from whole BM cells, LSK cells, or LK cells in 1% β-mercaptoethanol RLT buffer (Qiagen) using a RNeasy Micro Kit (Qiagen) according to the manufacturer’s instruction. Complementary DNA synthesis was performed using QuantiTect-Reverse -Transcription Kit (Qiagen). PCR amplifications were performed in three technical replicates for each gene of interest using Taqman Universal Master-Mix (Applied Biosystems) and measured using a QuantStudio-7 Flex Real-Time PCR System (Applied Biosystems). The expression level of messenger RNA for *Gata2* (Mm00492301, Applied Biosystems) or *Asxl1* (Mm01240150, Applied Biosystems) was normalised to the housekeeping gene *Hprt* mRNA (Mm00446968, Applied Biosystems) and determined using the 2^-ΔΔCT^ method of relative expression (52).

### RNA-Sequencing and analysis

Total RNA was isolated from purified HSCs (LSK CD48^-^CD150^+^) from 8-12 weeks old control, *Gata2^+/fl^; Vav-iCre^+^*, *Asxl1^+/fl^; Vav-iCre^+^*, or *Gata2^+/fl^, Asxl1^+/fl^, Vav-iCre^+^* mice using the RNeasy Plus Micro Kit (Qiagen) as described above. Assessment of total RNA quality and quantity was firstly performed using Agilent 2100 Bioanalyzer and RNA Nano 6000 kit (Agilent Technologies). According to the manufacturer’s instructions, RNA samples with a concentration of 0.5-6 ng and a RIN value of more than 8 were used for ribosomal RNA depletion utilizing a NEBNext rRNA Depletion Kit (E6310, New England Biolabs) and followed by library construction using using the NEB Ultra™ II Directional RNA Library Prep Kit for Illumina (New England Biolabs). Finally, constructed libraries were ultimately sequenced using a 75-base paired-end (2×75bp PE) dual index read format on the Illumina® HiSeq4000. RNA-Seq and bioinformatics were performed as described previously (8). The DEseq2 Bioconductor package in R-scripts statistical computing software was used to identify differentially expressed genes (DEGs) (53). DEGs with a cut-off of *P* < 0.05 and FDR < 0.05 were used for analysis of enriched biological pathways (54). The Ingenuity Pathway Analysis (IPA, Qiagen-Bioinformatics) software was utilized to explore gene ontology and common canonical pathways for DEGs. Gene Set Enrichment Analysis (GSEA) was also performed using hallmark gene-sets and C2-curated gene-sets (KEGG, Reactome, and BioCarta databases) in the GSEA Molecular Signatures Database (MSigDB, Version 4.0.2) (55). The Morpheus online software (Broad Institute) was utilized to generate heatmaps for differentially expressed genes.

### RNA-seq Data Availability

The accession number for the *Gata2^+/fl^; Vav-iCre^+^* HSC RNA-seq reported in this paper is GEO: GSE133248. The accession number for the *Gata2^+/fl^, Asxl1^+/fl^, Vav-iCre^+^* HSC RNA-seq reported in this paper will be published in due course.

### Statistical analysis

Figures were prepared by Prism 8.3 software (GraphPad Software, Inc). Statistical significance was determined by Mann–Whitney *U* test, Unpaired two tailed student’s *t* test, or One-way ANOVA - Tukey’s multiple comparisons test as follows: *, P < 0.05; **, P < 0.01; ***, P < 0.001; and ****, P <0.0001. For statistical analysis comparing two groups, all data underwent normality tests using Anderson-Darling and D’Agostino & Pearson tests. Data are presented as mean ± standard error of mean (SEM).

## Results

### Generation and validation of *Gata2^+/fl^; Vav-iCre^+^* haploinsufficient mice

To explore the function of *Gata2* haploinsufficiency in the adult hematopoietic system, *Gata2^+/+^; Vav-iCre^+^* with *Gata2^fl/fl^; Vav-iCre^-^* were crossed to produce *Gata2^+/fl^; Vav-iCre^+^* (heterozygote) and *Gata2^+/fl^; Vav-iCre^-^*(control) littermates (**Figure 1A**). *Gata2^+/fl^; Vav-iCre^+^* mice were born in the expected Mendelian ratio and appeared phenotypically normal when compared with their control littermates. To verify *Gata2* deletion, genotyping of ear notch biopsies and BM cells was conducted, which confirmed *Gata2* heterozygote or control status (**Supplementary Figure 1A**). Furthermore, *Gata2* mRNA expression was assessed in total BM cells and sorted LSK cells in *Gata2^+/fl^; Vav-iCre^+^* and control mice. Expression level of *Gata2* mRNA was similar in total BM in both littermates yet a two-fold reduction of *Gata2* mRNA was observed in HSPC enriched LSK cells in *Gata2^+/fl^; Vav-iCre^+^* when compared with control mice (**Supplementary Figure 1B**). Thus, *Gata2^+/fl^; Vav-iCre^+^* mice demonstrate efficient genetic deletion of a single *Gata2* allele with a reduction in *Gata2* expression in HSPCs.

**Figure 1.**
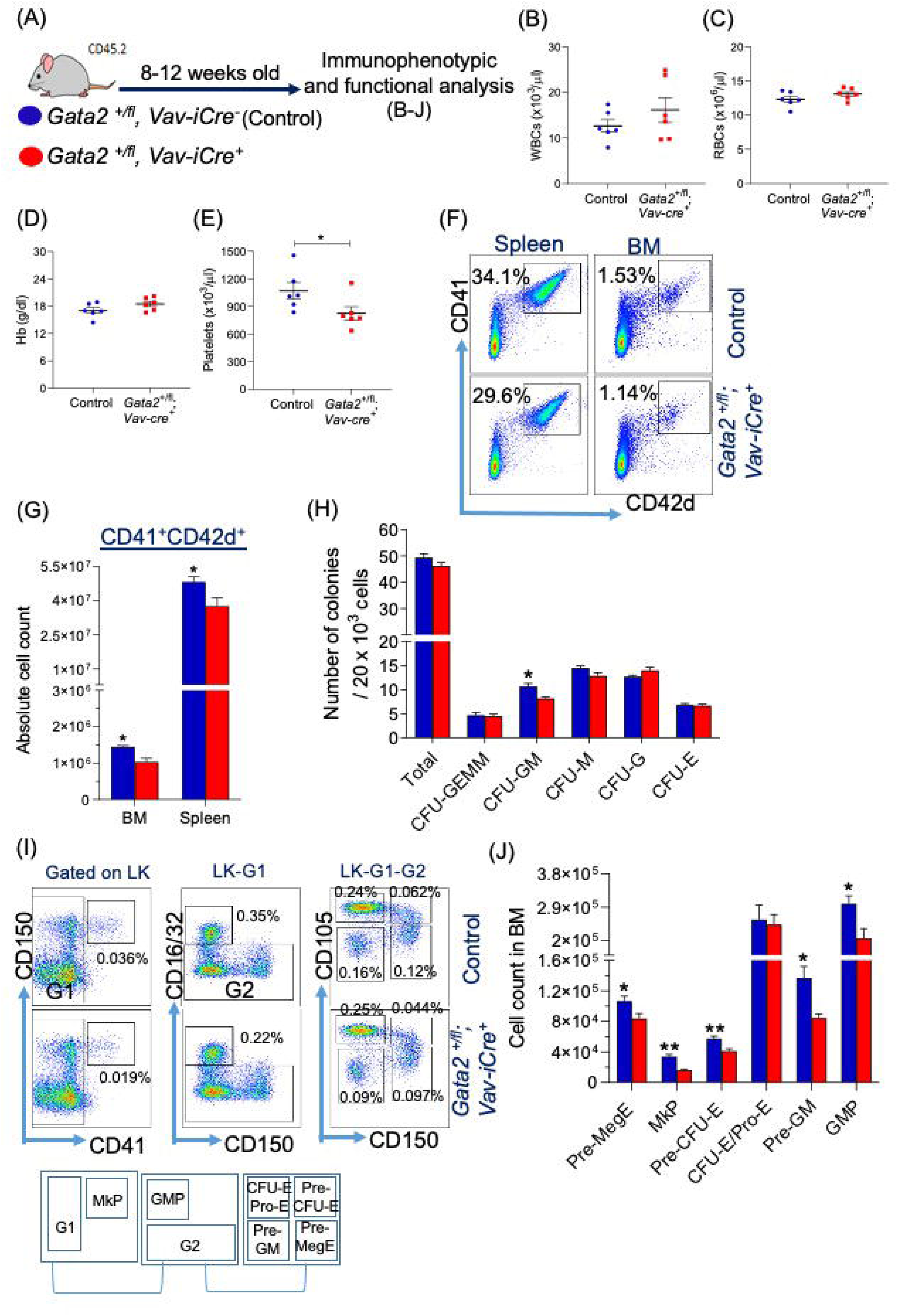
*Gata2* haploinsufficiency impairs megakaryopoiesis and committed myeloid progenitors. (A) Schematic depiction illustrates experimental design. *Gata2^+/fl^; Vav-iCre^−^* (control) and *Gata2^+/fl^; Vav-iCre^+^* littermates were assessed at 8-12 weeks old. (B, C, D, and E) Enumeration of mature peripheral blood WBCs (B), RBC (C), haemoglobin level (D), and platelets (E) in control and *Gata2^+/fl^; Vav-iCre^+^* mice. n= 6 per group from two independent experiments. (F and G) Representative flow cytometric analysis of the frequency of mature megakaryocytes (CD41^+^CD42d^+^) (F) and absolute numbers (G) in the BM and spleen from control and *Gata2^+/fl^; Vav-iCre^+^* mice. n= 4 mice per group from a single biological replicate. (H) Colony-forming unit (CFU) assay. A total of 20×10^3^ BM cells of *Gata2^+/fl^; Vav-Cre^+^* and control mice were plated in M3434 semisolid methylcellulose media containing SCF, IL-3, IL-6 and Epo, and formed colonies were counted 12 days after plating. n= 6 mice for each genotype from three independent experiments. Presented data are mean ± SEM. Statistical analysis: Mann-Whitney U test *, *P* < 0.05; **, *P* < 0.01. (I and J) Representative immunophenotypic analysis (I-Top) and gating strategy (I-Bottom) and absolute numbers (J) of BM of myeloid (Pre-GM, LKCD150^-^CD41^-^CD16/32^-^ CD105^-^ and GMPs, LK_CD150^-^_CD41^-^_CD16/32^+^), erythroid (Pre-MegE, LK_CD150^+^_CD41^-^_CD16/32^-^_CD105^-^; Pre-CFU-E, LK_CD150^+^_CD41^-^_CD16/32^-^_CD105^+^; and CFU-E/Pro-E, LK_CD150^-^_CD41^-^_CD16/32^-^_CD105^+^), and megakaryocyte (Pre-MegE and MkP, LK_CD150^+^_CD41^+^) progenitors from control (n=6) and *Gata2^+/fl^; Vav-iCre^+^* (n=6) mice from two independent experiments.

### *Gata2* haploinsufficiency causes defective early erythroid commitment and impairs platelet numbers mapping to an early defect in megakaryocyte progenitor generation

We initially evaluated the haematopoietic potential of *Gata2^+/fl^; Vav-iCre^+^* and control (*Gata2^+/fl^; Vav-Cre*^-^) mice by complete blood count (CBC) analysis. While white blood cells (WBCs), red blood cells (RBCs), and hemoglobin levels were equivalent between both genotypes, the number of platelets was significantly decreased in *Gata2^+/fl^;Vav-Cre^+^*mice when compared to control mice (**Figure 1B-E**), indicating that *Gata2* haploinsufficiency disrupts megakaryopoiesis in agreement with data implicating *Gata2* as a key regulator of megakaryopoiesis (6, 13). To uncover the cellular basis for the defect in platelet generation in *Gata2^+/fl^; Vav-Cre^+^* mice, we evaluated the frequency of mature CD41^+^CD42d^+^ megakaryocyte cells in BM and spleen, which were significantly reduced in BM and spleen in *Gata2^+/fl^; Vav-iCre^+^* (**Figure 1F** and **G**). Thus, *Gata2* haploinsufficiency perturbs megakaryocytic maturation.

To explore whether other lineage-specific haematopoietic cells in the PB, BM, spleen, and thymus were affected in *Gata2^+/fl^; Vav-Cre^+^* mice, flow cytometric analysis was performed for myeloid, erythroid, T and B-cell markers. We first assessed cellularity in hematopoietic organs in *Gata2^+/fl^; Vav-Cre^+^* mice and found that BM, spleen, and thymic cellularity was equivalent between genotypes (**Supplementary Figure 1C**). Furthermore, frequencies of fully differentiated myeloid, lymphoid, and erythroid cells in the PB, BM, spleen, and thymus were similar in both *Gata2^+/fl^; Vav-iCre^+^* and control mice (**Supplementary Figure 1D-G**). Functional analysis of BM myelo-erythroid lineage output in colony-forming cell (CFC) assays was comparable between genotypes, except for CFC-GM frequency, which was significantly reduced in *Gata2* haploinsufficient mice as previously demonstrated (11) (**Figure 1H**).

We next assessed the impact of *Gata2* haploinsufficiency on committed myeloid and lymphoid progenitors in BM. Committed myeloid restricted progenitor immunophenotyping has been defined by (49). In this paradigm of hematopoietic cell differentiation, common myeloid progenitors (CMPs) give rise to either primitive granulocyte/macrophage progenitors (Pre-GM, LK_CD150^-^_CD41^-^_CD16/32^-^_CD105^-^) that differentiate into granulocyte/macrophage progenitors (GMPs) (LK_CD150^-^_CD41^-^_CD16/32^+^) or primitive erythroid/megakaryocyte progenitors (Pre-MegE, LK_CD150^+^_CD41^-^_CD16/32^-^_CD105^-^). Pre-MegE cells segregate into either primitive erythroid forming colonies (Pre-CFU-E, LK_CD150^+^_CD41^-^_CD16/32^-^_CD105^+^) that further differentiate into erythroid forming colonies and pro-erythroblast (CFU-E and Pro-E, LK_CD150^-^_CD41^-^_CD16/32^-^_CD105^+^) or megakaryocyte progenitors (MkP, LK_CD150^+^_CD41^+^). Our analysis revealed that Pre-GM, GMP, Pre-MegE, MkP, and Pre-CFU-E were significantly reduced in *Gata2^+/fl^; Vav-iCre^+^* (**Figure 1I** and **J**). These data suggest that *Gata2* haploinsufficiency causes defects in early erythroid commitment. In contrast, common lymphoid progenitors, defined as Lin^-^_Sca1^low^_c-Kit^low^_CD127^+^ (47), were unchanged between the two genotypes (**Supplementary Figure 1H and I**). Early myeloid progenitor commitment in the BM, including megakaryocyte progenitor generation, was therefore compromised in the setting of *Gata2* haploinsufficiency. Together data suggest that reduced platelets observed in *Gata2^+/fl^; Vav-iCre^+^* mice arise from an early Pre-MegE and MkP progenitor defect and a subsequent block in differentiation toward maturing CD41^+^CD42d^+^ megakaryocytes.

### Reduced immunophenotypic HSPC abundance in *Gata2* haploinsufficient mice

To assess if the alteration in lineage-specific hematopoietic progenitors was caused by a disruption in the frequencies of BM HSPCs in *Gata2* haploinsufficient mice, we analysed the HSPC compartment by flow cytometry. LSK (Lin^-^_Sca-1^+^_c-Kit^+^) and SLAM markers (CD150 and CD48) were used to dissect HSPC compartments as follows: HSC, LSK_CD150^+^_CD48^-^; HPC1, LSK_CD150^-^_CD48^+^; HPC2, LSK_ CD150^+^_CD48^+^; and MPP, LSK_CD150^-^_CD48^-^ (46, 48) (**Figure 2A**). HPC1 progenitors differentiate into lymphoid/myeloid restricted progenitors and are incapable of advancing to megakaryocytes/ erythrocytes, whereas HPC2 cells are able to differentiate to all restricted hematopoietic progenitors (48). In parallel, HSCs give rise to lympho-myeloid progenitors (LMPPs), defined immunophenotypically as LSK_CD135^hi^_CD34^+^, that possess dual lymphoid/myeloid potential and overlap functionally with HPC1 (44). Frequencies of HSCs, MPPs, LMPPs and HPC1 were significantly reduced in BM from *Gata2^+/fl^; Vav-iCre^+^* compared to control mice, whereas the proportion of HPC2 was marginally increased in *Gata2^+/fl^; Vav-iCre^+^* as compared with control littermates, suggestive of a differentiation block at the transition from LMPP/HPC1 to HPC2 (**Figure 2B**).

**Figure 2.**
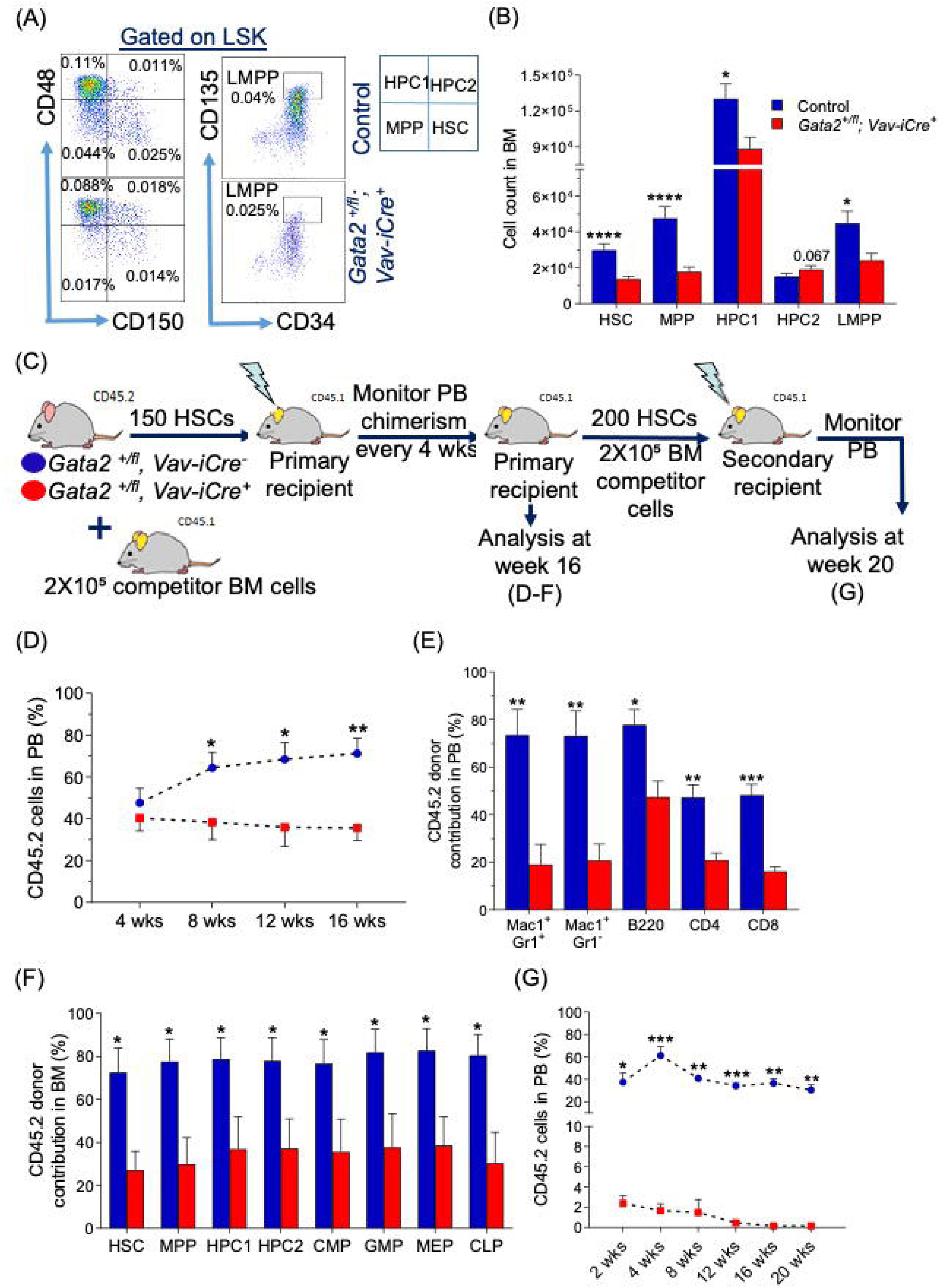
*Gata2* haploinsufficiency affects the development of primitive hematopoietic compartments and impairs multi-lineage reconstitution and self-renewal capacity of adult HSCs after transplantation. (A and B) Representative FACS plots (A-Left), gating strategy (A-Right), and absolute cell numbers of primitive HSPC compartments (B) including, HSCs (LSKCD150^+^CD48^−^), MPPs (LSKCD150^−^CD48^−^), HPC1 (LSKCD150^−^ CD48^+^), HPC2 (LSKCD150^+^CD48^−^), and LMPPs (LSKCD34^+^ CD135^hi^) in the BM from three or four independent experiments. n= 9 control and 10 *Gata2^+/fl^; Vav-iCre^+^* mice for HSCs, MPPs, HPC1, and HPC2, and n= 8 for each genotype for LMPPs. Statistical analysis: Unpaired, 2-tailed t-test for HSCs, MPPs, HPC1, and HPC2, and Mann-Whitney U test for LMPPs. (C) Experimental design of the competitive transplantation assay. For primary HSCs transplant, 150 CD45.2^+^ HSCs from either *Gata2^+/fl^; Vav-iCre^+^* or control mice alongside 2×10^5^ support CD45.1^+^ BM cells transplanted into lethally irradiated CD45.1^+^ recipients (n=8 mice for each group). Data were collected from three independent experiments. Five independent biological donors were used for each genotype. For secondary HSCs transplant, 200 CD45.2^+^ HSCs from either *Gata2^+/fl^; Vav-iCre^+^* or control derived cells from primary recipient mice were mixed with 2×10^5^ CD45.1^+^ competitor BM cells and transplanted into lethally irradiated secondary CD45.1^+^ hosts (n= 7 mice for each genotype). Data were collected from two independent experiments using 3-4 independent biological donors for each genotype. PB was assessed monthly for donor chimerism, and BM, spleen, and thymus cells were harvested and analysed at week 16 after transplantation. (D and E) Proportion of total CD45.2 cells at 4 weeks basis (D) and donor contribution (E) for myeloid (Mac1^+^Gr1^+^, Mac1^+^Gr1^-^), B-lymphoid (B220^+^), and T-lymphoid (CD4^+^, CD8^+^) in PB of primary transplanted recipients at week 16 post HSCs transplantation from control and *Gata2^+/fl^; Vav-iCre^+^* derived cells. Statistical analysis: Mann-Whitney U test. (F) Frequency of CD45.2 donor contribution to BM HSPCs (HSCs, MPPs, HPC1 and HPC2) and committed myeloid/lymphoid progenitors (CMPs, GMPs, MEPs, and CLPs) at week 16 after HSCs transplantation from control and *Gata2^+/fl^; Vav-iCre^+^* derived cells. Statistical analysis: Mann-Whitney U test. (G) Frequency of CD45.2 cells in PB at 2, 4, 8, 12, 16, 20 weeks after secondary HSC transplantation from primary recipients of control and *Gata2^+/fl^; Vav-Cre^+^* derived cells. Statistical analysis: Mann-Whitney U test. Data are presented as mean ± SEM. \**P* < 0.05, \*\**P* < 0.01, \*\*\**P* < 0.001, \*\*\*\**P* < 0.0001.

### *Gata2* haploinsufficient HSCs exhibit a functional defect in multi-lineage repopulation ability

Having shown that *Gata2* haploinsufficiency broadly reduces immunophenotypic HSPC abundance in BM, we assessed HSPC function *in vivo* using competitive transplantation experiments (50). Donor CD45.2 HSCs were prospectively isolated by Fluorescence-activated cell sorting (FACS) from either *Gata2^+/fl^; Vav-iCre^+^* or control mice and were mixed with competitor BM cells from CD45.1 mice and injected intravenously into CD45.1 irradiated recipient mice (**Figure 2C**). Overall, donor cell contribution from the *Gata2^+/fl^; Vav-iCre^+^* genotype to the PB was significantly decreased compared to the control group from week 8 post-transplant onwards (**Figure 2D**) and reflected multi-lineage engraftment defects in the PB, BM, spleen and thymus of recipients of *Gata2^+/fl^; Vav-iCre^+^*HSCs at week 16 post-transplantation (**Figure 2D** and **E** **and Supplementary Figure 2A – C**). To assess if *Gata2^+/fl^; Vav-iCre^+^* HSC repopulation defects stem from reduced engraftment defects of immunophenotypically defined HSPCs and committed progenitors, we immunophenotyped HSPC donor contribution, and found a significant reduction spanning all HSPCs populations and myeloid/lymphoid committed progenitors from *Gata2* haploinsufficient HSCs donor cells (**Figure 2F**). Thus, *Gata2* haploinsufficient HSCs lack competence for multi-lineage differentiation *in vivo* post transplantation.

### *Gata2* haploinsufficiency abolishes HSC self-renewal potential *in vivo*

Given that *Gata2* haploinsufficiency impairs HSC mediated multi-lineage differentiation post transplantation with a reduction in HSC engraftment (**Figure 2F**), we next assessed the impact of *Gata2* haploinsufficiency on HSC self-renewal, as judged by serial transplantation experiments (50). At week 16 of primary transplantation, FACS purified HSCs from *Gata2^+/fl^; Vav-iCre^+^* or control (CD45.2 cells) primary recipients were mixed with fresh competitor BM cells (CD45.1 cells) and subsequently re-transplanted to lethally irradiated secondary recipient mice (CD45.1 background) (**Figure 2C**). Donor chimerism in PB of secondary recipients receiving *Gata2^+/fl^; Vav-iCre^+^* HSCs was severely reduced from week 2 post-transplant when compared to the control group and, remarkably, was nearly undetectable by week 20 post-transplantation in both PB and BM (**Figure 2G** **and Supplementary Figure 2D**). These data indicate that *Gata2* haploinsufficiency abolishes HSC self-renewal *in vivo*.

### *Gata2* haploinsufficient HSCs display disrupted B-cell development, DNA damage responses and inflammatory signalling transcriptional networks

To investigate the molecular underpinnings of *Gata2* haploinsufficiency induced defects in HSC function, we conducted genome wide transcriptomic analysis by RNA-Seq of highly purified HSCs (LSK_CD150^+^CD48^-^) from *Gata2^+/fl^; Vav-iCre^+^* and control mice. 117 statistically significant differentially expressed genes (DEGs) were identified in *Gata2* haploinsufficient HSCs relative to control HSCs, of which 74 genes were down-regulated and 43 genes were up-regulated (**Supplementary Figure 3A**).

Ingenuity Pathway Analysis (IPA) of differentially regulated genes from *Gata2* haploinsufficient HSCs indicated molecular function affected by *Gata2* in normal hematopoiesis (e.g. cell death and apoptosis/survival, cellular growth and proliferation, hematological system development and function) (**Supplementary 3B and C**). Notably, cell-to-cell signalling and interaction, as evidenced by deregulated cytoskeletal organization, and cell adhesion genes (**Supplementary Figure 3B and C**), both of which are required for maintaining HSC integrity (56–58), were also differentially deregulated in *Gata2* haploinsufficient HSCs. Pathways directly related to *GATA2* clinical haploinsufficiency syndromes were observed in *Gata2* haploinsufficient HSCs, including B-cell development/humoral responses (37–39). To validate the impact of this transcriptional signature on the B-cell developmental program in *Gata2* haploinsufficient mice, we analyzed B-cell progenitor development and found a selective expansion in early pre-B cells (IgM-CD43+CD19+BP1+) in BM (**Supplementary Figure 3D and E**) and a specific decrease in Pre-Pro B cells (IgM-CD43+CD19-BP1-) in the spleen of *Gata2* haploinsufficient mice (**data not shown**).

By analysis of Ingenuity Canonical Pathway (ICP), Gene Set Enrichment Analysis (GSEA) and Molecular Signatures Database (MSigDB, hallmark gene-sets and C2-curated gene-sets), we found *Gata2* haploinsufficient HSCs displayed altered DNA repair pathways, specifically double-stranded (ds) DNA-break repair by homologous recombination and base-excision repair (BER), and pro-inflammatory signalling (e.g. Toll-like receptor signalling, IFNγ signalling, IL-10 signalling, and granulocyte adhesion/diapedesis) as well as extracellular matrix components pathways implicated in maintenance of HSC integrity (e.g. glycosaminoglycan-protein biosynthesis and inhibition of matrix metalloproteases) (59, 60) (Figure 3A-C, **Supplementary Figure 3F**).

**Figure 3.**
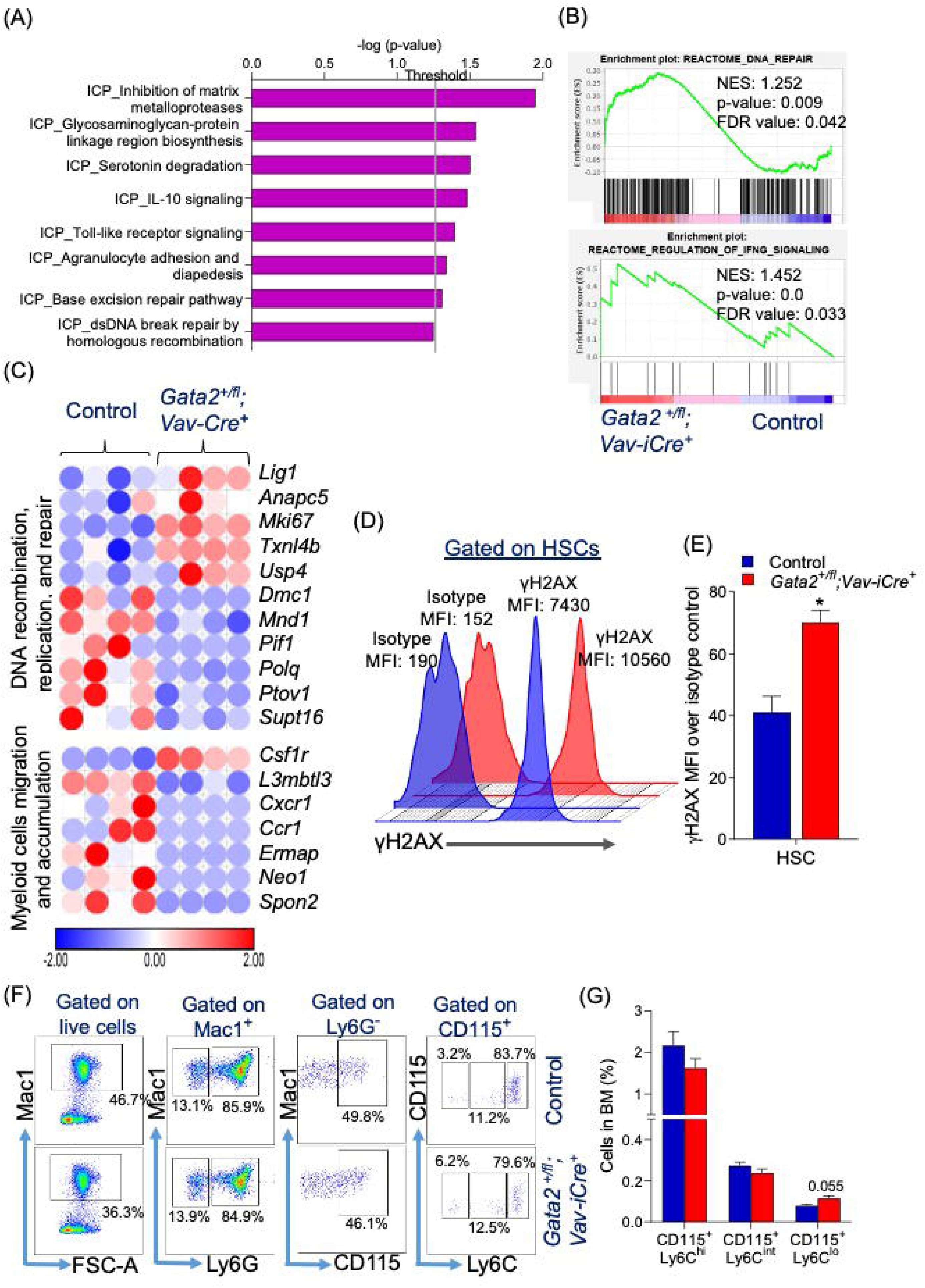
*Gata2* haploinsufficient HSCs display deregulation of transcriptional signatures associated with the DNA damage repair and proinflammatory signalling. RNA-Sequencing (RNA-Seq) of purified HSCs (LSKCD150^+^CD48^-^) from both control (n= 4) and *Gata2^+/fl^; Vav-iCre^+^* (n= 4) mice. (A) Biological pathway analysis of differentially expressed genes in *Gata2* haploinsufficient HSCs relative to control HSCs using IPA software. Given data are presented as –log_10_ (*P* value), and the threshold in the grey colour indicates a *P* value of 0.05. ICP indicates Ingenuity Canonical Pathway. Enriched pathways are determined by Fischer’s Exact Test. (B) GSEA plots display enriched pathways for up-regulated genes in *Gata2* haploinsufficient HSCs compared with control HSCs. Statistical significance is determined by p-values and FDR-values at <0.05 level. NES indicates normalised enrichment score. (C) Heatmaps of the differentially significant genes in *Gata2* haploinsufficient HSCs related to control HSCs. Heatmaps are drawn by Morpheus online software showing a z-score scale. The red colour indicates up-regulated genes; and blue colour, down-regulated genes. The colour gradient represents the intensity of gene expression. (D and E) Representative histogram plot analysis (D) exhibiting the median fluorescence intensity (MFI) of γH2AX expression and bar chart analysis (E) of the ratio of γH2AX MFI with respect to fluorescent isotype-control in BM HSCs from control (n= 4) and *Gata2^+/fl^; Vav-iCre^+^* (n= 4) mice from two independent experiments. Presented data are mean ± SEM. Statistical analysis: Mann-Whitney U test *, P < 0.05. (F and G) Representative immunophenotypic analysis (F) and proportions (G) of BM phagocytic and pro-inflammatory + macrophages (Mac1^+^ Ly6G^−^ CD115^+^ Ly6C^high^), pro-inflammatory macrophages (Mac1^+^ Ly6G^−^ CD115^+^ Ly6C^intermediate^) and anti-inflammatory macrophages (Mac1^+^ Ly6G^−^ CD115^+^ Ly6C^low^) from control (n= 5) and *Gata2^+/fl^; Vav-iCre*^+^ (n= 5) mice from two independent experiments. Data are presented as mean ± SEM. Statistical analysis: Mann-Whitney U test.

To validate our observation of altered transcriptional programming in DNA damage response pathways at the protein level in *Gata2* haploinsufficient HSCs, we used flow cytometry-based analysis of γH2AX, a member of the histone H2A family that becomes phosphorylated at Ser139 following dsDNA break damage and accumulates at locations of DNA damage to attract DNA-damage response proteins (61). γH2AX expression, as gauged by median fluorescence intensity (MFI), was increased in BM HSCs in *Gata2^+/fl^; Vav-iCre^+^* (**Figure 3D and E**), supporting the notion that *Gata2* haploinsufficiency induces dsDNA break damage in HSCs.

As IPA identified deregulated GATA2 mediated IFNγ target genes and cell adhesion molecules in HSCs (**Figure 3B**), both of which are involved in the recruitment of macrophages that modulate inflammation (**Figure 3B** and **C**), we sought to assess the impact of altered inflammatory signalling in *Gata2* haploinsufficient HSCs on the generation of downstream inflammatory cell subsets in BM and spleen. By using flow cytometry, we evaluated functionally distinct Mac1^+^ Ly6G^−^ CD115^+^ macrophage populations by Ly6C expression that discriminate between phagocytic and pro-inflammatory macrophages (Mac1^+^ Ly6G^−^ CD115^+^ Ly6C^high^), pro-inflammatory macrophages (Mac1^+^ Ly6G^−^ CD115^+^ Ly6C^intermediate^) and anti-inflammatory macrophages (Mac1^+^ Ly6G^−^ CD115^+^ Ly6C^low^) (62). While pro-inflammatory macrophage populations were unchanged in *Gata2* haploinsufficient mice, anti-inflammatory macrophages were marginally increased within the BM of *Gata2* haploinsufficient mice (**Figure 3F and G**). Phagocytic and pro-inflammatory macrophages were decreased in the spleen of *Gata2* haploinsufficient mice while splenic anti-inflammatory macrophages were increased **(Supplementary Figure 3G)**. In agreement with attenuated inflammatory responses to infectious stimuli in *Gata2* haploinsufficient mice (63), these data suggest altered inflammatory signalling in *Gata2* haploinsufficient HSCs reduces the generation of inflammatory populations in BM and spleen.

### *Ex vivo* knockdown of *Asxl1* in Gata2 haploinsufficient BM cells reduces multi-potent hematopoietic progenitor generation

Next, we sought to explore potential genetic pathways contributing to impaired DNA damage and inflammatory responses in *Gata2* haploinsufficient mice by assessing the role of *Asxl1*, an epigenetic modifier involved in the regulation of Polycomb genes in normal hematopoiesis and hematological neoplasms (41, 64). *ASXL1* deregulation has been implicated in DNA damage accrual in HSCs (65), and somatic *ASXL1* mutations are a frequent abnormality observed in clonal hematopoiesis of indeterminate potential (CHIP) that drives deregulated inflammation and the propensity to develop hematologic malignancies during aging (41, 64, 65). We also elected to study *Asxl1* as acquired *ASXL1* mutations have been detected in approximately 30% of MDS/AML cases with germline *GATA2* haploinsufficiency mutations (33, 34).

To mimic *GATA2* haploinsufficiency with acquired *ASXL1* somatic mutations and explore the interaction between GATA2 and ASXL1 pathways in HSPCs, we used shRNA mediated knockdown of *Asxl1* in *Gata2^+/fl^; Vav-iCre^+^* or control (WT) BM HPSCs (LSK cells) to functionally analyse the impact of *Gata2/Asxl1* deficiency on hematopoietic progenitor function (**Figure 4A**). Using a lentiviral vector linking a mCherry reporter to *Asxl1* shRNAs (*Asxl1*-sh1 and *Asxl1*-sh2) or empty vector (EV) control, we initially validated knockdown of *Asxl1* in WT LKS cells transduced with each vector (**Figure 4A**). Equivalent transduction efficiency, nearing 60%, was observed in WT LKS cells transduced with EV and *Asxl1* shRNA groups (**Figure 4B**). Furthermore, *Asxl1*-sh1 and *Asxl1*-sh2 displayed an approximately twofold and fourfold reduction in *Asxl1*-mRNA respectively, compared to EV transduced WT LKS cells (**Figure 4C**). After transduction of LSK cells harvested from WT or *Gata2^+/fl^; Vav-iCre^+^* mice with EV or *Asxl1-sh1 and Asxl1-sh2,* mCherry^+^ cells were isolated by FACS and plated in a colony forming cell assay (CFC) assay, where total CFC number was significantly decreased in all experimental groups when compared to EV transduced WT LKS cells (**Figure 4D**). As expected, total CFC number was reduced in WT LKS cells transduced with *Asxl1-sh1 and Asxl1-sh2* and, to a lesser extent, in single haploinsufficient *Gata2* transduced with EV (**Figure 4D**). Notably, there was a significant reduction in total CFCs in the *Gata2^+/fl^; Vav-iCre^+^*-*Asxl1*-sh1/sh2 group compared to single haploinsufficient *Gata2* transduced with EV or WT LKS cells transduced with *Asxl1* shRNA (**Figure 4D**). This reduction in CFCs from *Gata2^+/fl^; Vav-iCre^+^*-*Asxl1*-sh1/sh2 LKS cells reflected a substantial reduction across multipotential progenitors (CFU-GEMM) and bi-potent myeloid progenitors (CFC-GM) compared to single haploinsufficient *Gata2* transduced with EV or WT LKS cells transduced with *Asxl1* shRNA (**Figure 4E**). All lineage specific progenitors (CFU-GM, CFU-M, CFU-G, and CFU-E) were reduced in *Gata2^+/fl^; Vav-iCre^+^*-*Asxl1*-sh1/sh2 LKS cells compared to single haploinsufficient *Gata2* transduced with EV (**Figure 4E**). Thus, knockdown of *Asxl1* in *Gata2* haploinsufficient LSK cells functionally blocks the differentiation capacity of multipotent progenitors *in vitro*. These data further suggest knockdown of *Asxl1* potentiates a defective differentiation program already established in *Gata2* haploinsufficient mice (**Figure 1** **and Supplementary Figure 1**) and observed following transplantation of *Gata2* haploinsufficient HSCs (**Figure 2** **and Supplementary Figure 2**).

**Figure 4.**
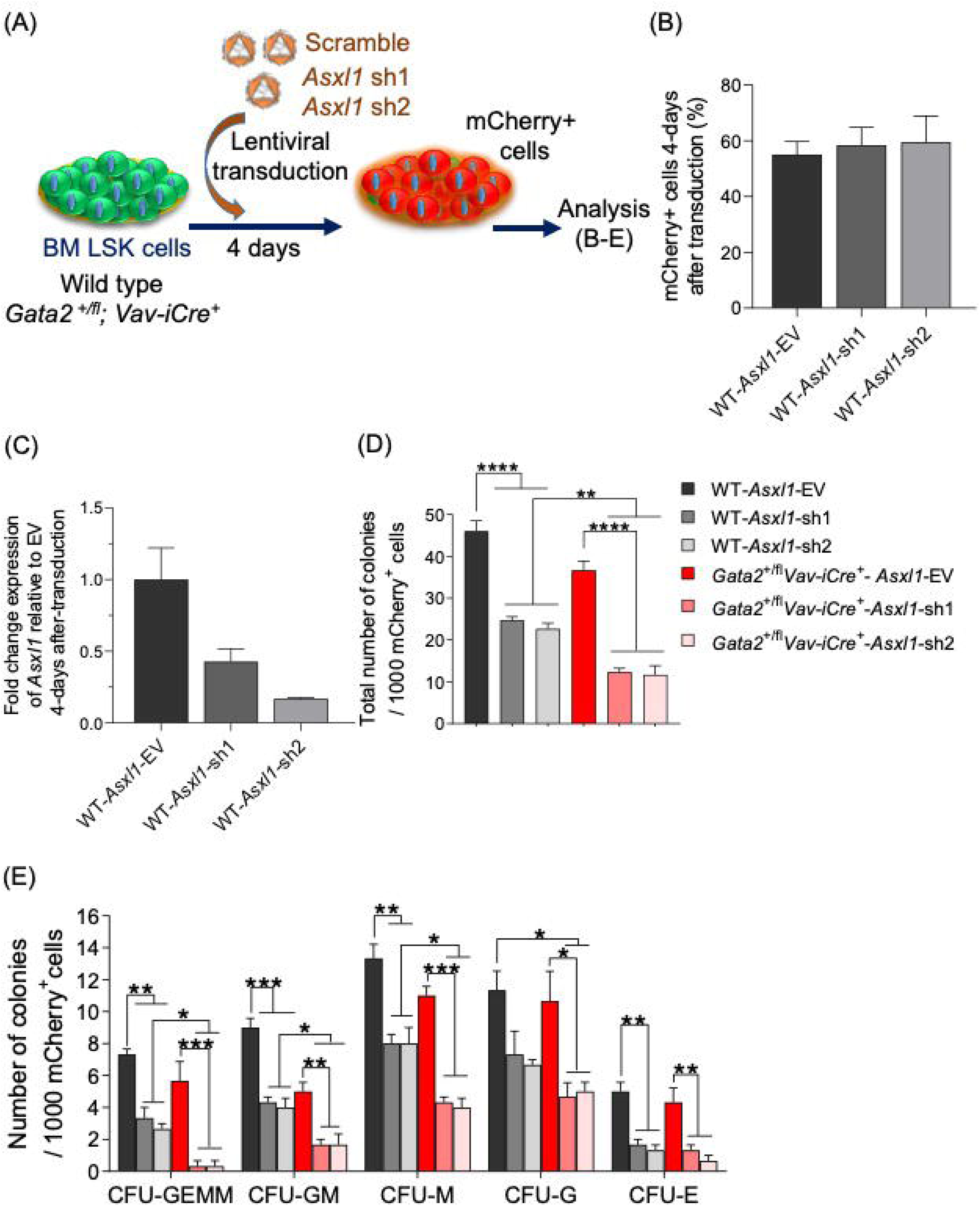
*In vitro* knockdown of *Asxl1* in *Gata2* haploinsufficient BM HSPCs impairs their differentiation capacity into myeloid progenitors. (A) Schematic depiction of *in vitro Asxl1* knockdown experiment. A total of 50×10^3^ freshly sorted LSK cells from either wild-type or *Gata2^+/fl^; Vav-iCre^+^* mice were transduced with *Asxl1-*EV, *Asxl1-*sh1, or *Asxl1-*sh2 at MOI of 50 and incubated overnight in IMDM media supplemented with 10% FBS, 1% L-glutamine, 1% Penicillin/Streptomycin, 100 ng/ml m-SCF, 100 ng/ml m-TPO, and 100 ng/ml m-Flt3L. Four days post transduction, mCherry^+^ cells were evaluated by FACS and collected by FACS-sorter for qPCR analysis and CFC assays. (B) The frequency of mCherry^+^ cells 4-days after transduction in transduced wild-type LSK cells (n= 3) with EV and *Asxl1* hairpins from two independent experiments. (C) The fold-change expression level of *Asxl1* mRNA in transduced wild-type LSK cells (n=2) relative to *Asxl1-*EV from two independent experiments. (D and E) Total colony numbers (D) and differentiated myeloid colonies (E) 10-12 days post plating in methylcellulose media of sorted mCherry^+^ BM cells from WT-*Asxl1*-EV (control), WT-*Asxl1*-sh1, WT-*Asxl1*-sh2, *Gata2^+/fl^; Vav-iCre* -*Asxl1*-EV, *Gata2^+/fl^; Vav-iCre* -*Asxl1*-sh1, *Gata2^+/fl^; Vav-iCre* - *Asxl1*-sh2 transduced cells. n= 3 per genotype from 2 independent experiments. Data are presented as mean ± SEM. Statistical analysis: One-Way ANOVA with Tukey’s multiple comparisons test *, P < 0.05; **, P < 0.01; ***, P < 0.001.

### Immunophenotypic HSPC and progenitor defects in *Gata2* and *Asxl1* double haploinsufficient mice mirror those found in *Gata2* haploinsufficient mice

That knockdown of *Asxl1* in *Gata2* haploinsufficient LKS cells results in a reduction of multipotent progenitors beyond that observed in single haploinsufficient *Gata2* or *Asxl1* LKS cells, suggests genetic interaction between GATA2➔ASXL1 pathways in HSPCs (8, 66). To formally test this *in vivo*, we assessed whether combined *Gata2* and *Asxl1* haploinsufficiency causes exacerbated loss of HSPCs when compared to either *Gata2* or *Asxl1* haploinsufficiency. To generate compound *Gata2/Asxl1* haploinsufficient mice, *Gata2^+/fl^, Asxl1^+/fl^, Vav-iCre^-^* males were bred with *Gata2^+/+^; Asxl1^+/+^; Vav-iCre^+^* females to generate *Gata2^+/fl^; Asxl1^+/fl^; Vav-iCre^+^* (double *Gata2/Asxl1* heterozygote mice), *Gata2^+/fl^; Asxl1^+/fl^; Vav-iCre^-^* (control mice), *Gata2^+/fl^; Vav-iCre^+^* (single *Gata2* heterozygote), and *Asxl1^+/fl^; Vav-iCre^+^* (single *Asxl1* heterozygote) (**Figure 5A**). All littermates were born healthy, in the predicted Mendelian proportions. Genotyping PCR of ear notch biopsies and BM cells confirmed complete excision of one allele of both *Gata2* and *Asxl1* genes (**Supplementary Figure 4A**) and mRNA analysis confirmed reduced *Asxl1* expression in total BM cells from *Asxl1^+/fl^; Vav-iCre^+^* and a decrease in *Asxl1* and *Gata2* mRNAs in HSPC enriched c-kit^+^ cells from *Gata2^+/fl^; Asxl1^+/fl^; Vav-iCre^+^* mice (**Supplementary Figure 4B and C**).

**Figure 5.**
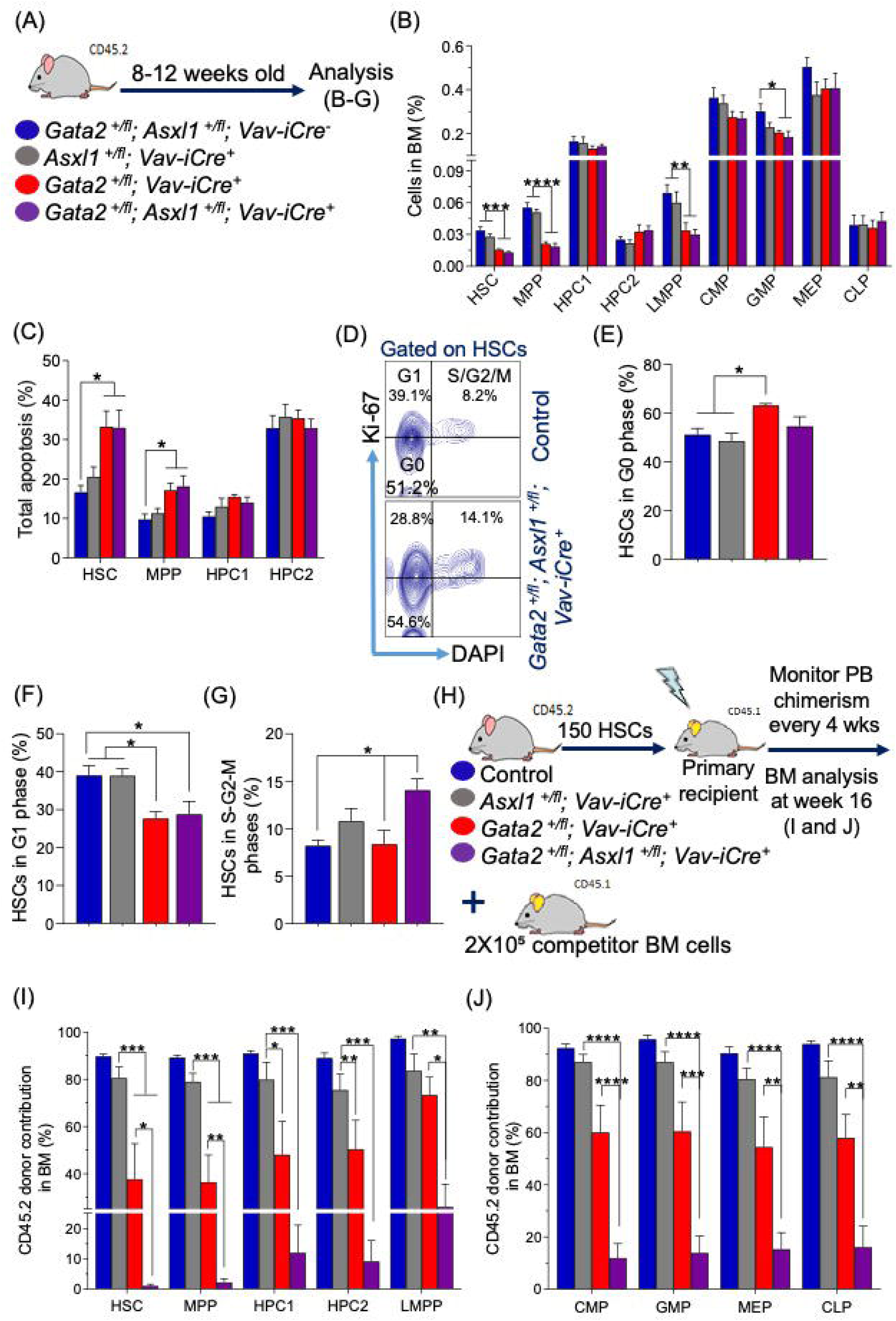
*Gata2*/*Asxl1* double haploinsufficient mice demonstrated increased HSCs cycling, decreased HSCs survival, and ultimately exhausted HSC pool size after HSCs transplantation. (A) Experimental design for double haploinsufficient mice of *Gata2* and *Asxl1*. *Gata2^+/fl^; Asxl1^+/fl^; Vav-iCre^-^* (Control), *Asxl1^+/fl^; Vav-iCre^+^*, *Gata2^+/fl^; Vav-iCre^+^*, and *Gata2^+/fl^; Asxl1^+/fl^; Vav-iCre^+^* mice were assessed at 8-12 weeks of age. (B) Frequency of BM HSPCs (HSCs, MPPs, HPC1, HPC2, and LMPPs) and committed myeloid/lymphoid progenitors (CMPs, GMPs, MEPs, and CLPs) from control (n=6), *Asxl1^+/fl^; Vav-iCre^+^* (n=5), *Gata2^+/fl^; Vav-iCre^+^* (n=6), and *Gata2^+/fl^; Asxl1^+/fl^; Vav-iCre^+^* (n=6) mice from two independent experiments. (C) Proportion of total apoptosis in BM HSPCs (HSCs, MPPs, HPC1, and HPC2) from control (n=6), *Asxl1^+/fl^; Vav-iCre^+^* (n=5), *Gata2^+/fl^; Vav-iCre^+^* (n=6), and *Gata2^+/fl^; Asxl1^+/fl^; Vav-iCre^+^* (n=6) mice from two independent experiments. The apoptosis assay was carried out using annexin V and DAPI stains. (D, E, F, and G) Representative FACS plots of the cell cycle (D) and frequencies of BM HSPCs (HSCs, MPPs, HPC1, and HPC2) in the G0 phase (E), G1 phase (F), and S/G2/M phases (G) from control (n=4), *Asxl1^+/fl^; Vav-iCre^+^* (n=4), *Gata2^+/fl^; Vav-iCre^+^* (n=4), and *Gata2^+/fl^; Asxl1^+/fl^; Vav-iCre^+^* (n=4) mice from two independent experiments. The cell cycle status was assessed using Ki-67 and DAPI stains, in which G0 indicates Ki-67^-^DAPI^-^; G1, Ki-67^+^DAPI^-^; and S/G2/M, Ki-67^+^DAPI^+^. (H) Schematic depiction of the competitive HSC transplantation. Lethally irradiated CD45.1^+^ recipients were injected with 150 CD45.2^+^ HSCs from control, *Gata2^+/fl^; Asxl1^+/fl^; Vav-iCre^+^*, *Gata2^+/fl^; Vav-iCre^+^*, and *Gata2^+/fl^; Asxl1^+/fl^; Vav-iCre^+^* mice together with 2×10^5^ competitor CD45.1^+^ BM cells. Four independent donor mice were used for each group. (I and J) Frequency of CD45.2 donor contribution at 16 weeks post transplantation in BM HSPCs (HSCs, MPPs, HPC1, HPC2, LMPPs) (I) and committed haematopoietic progenitors (CMPs, GMPs, MEPs, CLPs) (J) from control (n=7), *Asxl1^+/fl^; Vav-iCre^+^* (n=6), *Gata2^+/fl^; Vav-iCre^+^* (n=6), and *Gata2^+/fl^; Asxl1^+/fl^; Vav-iCre^+^* (n=6) donor derived-cells from two independent experiments. Presented data are mean ± SEM. Statistical analysis: One-Way ANOVA with Tukey’s multiple comparisons test *, P < 0.05; **, P < 0.01; ***, P < 0.001; ****, P < 0.0001.

Fully differentiated hematopoietic cells from PB, BM and spleen and cellularity of BM and spleen from young adult (8–12-week-old) *Gata2^+/fl^; Asxl1^+/fl^; Vav-iCre^+^* mice mirrored hematopoietic potential in their single heterozygote counterparts, indicating haploinsufficiency of both *Gata2* and *Asxl1* does not disrupt steady-state terminal differentiation of hematopoietic cells in PB, BM and spleen nor the cellularity of BM and spleen (**Supplementary Figure 4D-H**). Yet, the frequencies of HSCs, MPPs, LMPPs and GMP in the BM of double haploinsufficient mice was reduced compared to *Asxl1^+/fl^; Vav-iCre^+^* and control littermates and, notably, mirrored the HSPC/progenitor defects found in *Gata2^+/fl^; Vav-iCre^+^* mice (**Figure 5A and B**). No differences were observed in CMP, MEP, and CLP frequency comparing experimental genotypes to *Gata2^+/fl^; Asxl1^+/fl^; Vav-iCre^-^* controls (**Figure 5B**). Thus, *Gata2*/*Asxl1* double haploinsufficiency *in vivo* impacts immunophenotypic HSPCs/progenitors in a manner closely resembling that observed in *Gata2* haploinsufficient mice.

### Increased HSC proliferation potential in *Gata2* and *Asxl1* double haploinsufficient mice

While double *Gata2* and *Asxl1* haploinsufficiency replicated the HSPC immunophenotype found in *Gata2* haploinsufficient mice, it was unclear whether cellular mechanisms regulating HSPC genomic integrity, namely apoptosis and cell cycle status, were similar in both genotypes. Annexin V analysis revealed a significant increase in total apoptosis from HSCs and MPPs in *Gata2^+/fl^; Asxl1^+/fl^; Vav-iCre^+^* mice when compared to *Asxl1^+/fl^; Vav-iCre^+^* or control littermates, which, importantly, correlated with enhanced HSPC apoptosis observed in *Gata2^+/fl^; Vav-iCre^+^* mice (**Figure 5C**). Determination of HSC cell cycle status revealed an unchanged proportion of quiescent HSCs, and an enhancement in S/G2/M phase HSCs from *Gata2^+/fl^; Asxl1^+/fl^; Vav-iCre^+^* mice compared to *Gata2^+/fl^; Vav-iCre^+^* mice or control littermates (**Figure 5D**, **E and G**). The proportion of G1 HSCs was similar in *Gata2^+/fl^; Asxl1^+/fl^; Vav-iCre^+^* and *Gata2^+/fl^; Vav-iCre^+^* mice and reduced compared to *Asxl1^+/fl^; Vav-iCre^+^* or control littermates (**Figure 5D and F**). Thus, *Gata2* and *Asxl1* haploinsufficiency together cause an increase in HSC proliferation, contrasting sharply with an increase in HSC quiescence observed from *Gata2^+/fl^; Vav-iCre^+^* mice (**Figure 5D and E**). These data further suggest that combined *Gata2* and *Asxl1* haploinsufficiency compromises both unique and complementary mechanisms that regulate genomic integrity in the HSC pool.

### Functional exhaustion of HSCs following competitive transplantation of HSCs from *Gata2* and *Asxl1* double haploinsufficient mice

Having shown that *Gata2/Asxl1* double haploinsufficient HSCs were more proliferative, we evaluated how this would impact *Gata2^+/fl^; Asxl1^+/fl^; Vav-iCre^+^* HSC function under conditions of proliferative stress using competitive transplantation experiments. We transplanted HSCs from control, *Asxl1^+/fl^; Vav-iCre^+^*, *Gata2^+/fl^; Vav-iCre^+^*, or *Gata2^+/fl^; Asxl1^+/fl^; Vav-iCre^+^* CD45.2 mice alongside 2×10^5^ CD45.1 competitor BM cells into irradiated CD45.1-recipient mice, monitoring PB engraftment every 4 weeks and chimerism in BM and spleen of recipients at week 16 following transplantation (**Figure 5H**).

While equivalent PB engraftment was noted initially for all genotypes at week 4 post-transplant, from week 8 onwards, recipients of *Gata2/Asxl1* double haploinsufficient HSCs experienced a progressive decrease in PB chimerism that was statistically significant compared to recipients of control or *Asxl1^+/fl^; Vav-iCre^+^* HSCs (**Supplementary Figure 5A)**. In comparison to their control or *Asxl1^+/fl^; Vav-iCre^+^* counterparts, recipients of *Gata2/Asxl1* double haploinsufficient HSCs were significantly reduced in PB myeloid (Mac-1^+^Gr-1^+^ and Mac-1^+^Gr1^-^) engraftment capacity at week 16 (**Supplementary Figure 5B)**. As expected, recipients of HSCs from *Gata2^+/fl^; Vav-iCre^+^* displayed a significant reduction in PB engraftment from week 8 onwards when compared to mice receiving control HSCs (**Figure 2D** **and Supplementary Figure 5A**); at week 16 this led to diminishment of all lympho-myeloid engraftment in PB (**Supplementary Figure 5B**). In contrast, PB chimerism in recipients of *Asxl1^+/fl^; Vav-iCre^+^* HSCs was more moderately reduced over time compared to recipients of control HSCs (**Supplementary Figure 5A**). At 16 weeks post-transplant, this was reflected by a significant reduction in only T-cell engraftment to the PB of mice transplanted with HSCs from the *Asxl1^+/fl^; Vav-iCre^+^* genotype (**Supplementary Figure 5B**). Overall, *Gata2/Asxl1* double haploinsufficient HSCs were severely compromised in their PB engraftment capacity after transplantation. Notably, recipients of *Gata2/Asxl1* double haploinsufficient HSCs exhibited lower PB engraftment than recipients of either *Asxl1^+/fl^; Vav-iCre^+^* or *Gata2^+/fl^; Vav-iCre^+^*HSCs starting from week 8, though this reduction did not reach statistical significance compared to recipients receiving *Gata2^+/fl^; Vav-iCre^+^* HSCs (**Supplementary Figure 5A**).

To evaluate the provenance of defective PB engraftment in transplant recipients of *Gata2/Asxl1* double haploinsufficient HSCs, we assessed donor HSPCs and committed hematopoietic population contribution to the BM of recipients at week 16. Remarkably, HSCs and MPPs were barely detectable in *Gata2^+/fl^; Asxl1^+/fl^; Vav-iCre^+^* genotype recipients compared to either control, *Asxl1^+/fl^; Vav-iCre^+^*, or *Gata2^+/fl^; Vav-iCre^+^* recipients (**Figure 5I**). Other HSPCs (HPC1, HPC2, and LMPPs) and committed progenitors (CMPs, GMPs, MEPs, and CLPs) donor cell engraftment were markedly attenuated in *Gata2^+/fl^; Asxl1^+/fl^; Vav-iCre^+^* recipients compared to control or *Asxl1^+/fl^; Vav-iCre^+^* genotype recipients (**Figure 5I** and **J**). A statistically significant reduction in LMPP, CMP, GMP, MEP, and CLP donor cell engraftment was also noted in *Gata2^+/fl^; Asxl1^+/fl^; Vav-iCre^+^* recipients in comparison to *Gata2^+/fl^; Vav-iCre^+^* recipients (**Figure 5I** and **J**). As predicted, the frequency of HSPCs and hematopoietic progenitor donor cells from recipients of HSCs from *Gata2^+/fl^; Vav-iCre^+^* mice were significantly lower than their control counterparts, whereas recipients of *Asxl1^+/fl^; Vav-iCre^+^* HSCs showed comparable HSPC and committed progenitor repopulation potential to recipients of control HSCs (**Figure 5I** and **J**). Thus, engraftment of HSPC compartments to the BM of recipients from *Gata2^+/fl^; Asxl1^+/fl^; Vav-iCre^+^* was markedly reduced when compared to their single haploinsufficient counterparts, indicating a failure of *Gata2/Asxl1* double haploinsufficient HSCs to adequately reconstitute HSPC compartments that leads to a functional PB engraftment defect after transplantation.

### RNA-seq analysis of *Gata2/Asxl1* double haploinsufficient HSCs identifies common *Gata2* and *Asxl1* target genes/pathways and a unique transcriptomic signature

*Gata2*/*Asxl1* haploinsufficiency augments HSC proliferation and causes HSPC functional defects beyond that observed in their single haploinsufficient counterparts, suggesting cooperative genetic interaction between *Gata2* and *Asxl1* pathways. To explore the transcriptional programmes underpinning genetic interaction between *Gata2* and *Asxl1* pathways, we conducted RNA-Seq analysis of purified BM HSCs from control, *Asxl1^+/fl^; Vav-iCre^+^*, *Gata2^+/fl^; Vav-iCre^+^*, and *Gata2^+/fl^; Asxl1^+/fl^; Vav-iCre^+^* mice.

1251 down-regulated genes and 1355 up-regulated genes were found in double *Gata2*/*Asxl1* haploinsufficient HSCs, tallying up to 2606 differentially expressed genes (DEGs) in *Gata2*/*Asxl1* double haploinsufficient HSCs when compared to control HSCs (**Supplementary Figure 6A**). 2083 down-regulated genes and 2135 up-regulated genes, a total of 4218 DEGs, were identified in HSCs from *Asxl1^+/fl^; Vav-iCre^+^*mice in comparison to control HSCs (**Supplementary Figure 6B**). In agreement with the notion that *ASXL1* mutations act as a preleukemic initiator (33, 41, 67, 68), several affected pathways that are involved in HSC integrity were detected in *Asxl1* haploinsufficient HSCs, including DNA damage repair, cellular survival, metabolic processes, and AML signalling (**Supplementary Figure 6D**). Approximately 50% (2139) of DEGs were shared between *Gata2/Asxl1* double haploinsufficient HSCs and *Asxl1^+/fl^; Vav-iCre^+^* HSCs. Of 117 DEGs identified from *Gata2^+/fl^; Vav-iCre^+^* HSCs (**Supplementary Figure 6C**), approximately 39% (46 DEGs) were shared with *Gata2/Asxl1* double haploinsufficient HSCs (**Supplementary Figure 6C**). 39 DEGs were common to *Gata2* and *Asxl1* single haploinsufficient HSCs and *Gata2* and *Asxl1* double haploinsufficient HSCs (**Supplementary Figure 6C**). Together, these data demonstrate a core common transcriptional signature specifically relating to GATA2 and ASXL1 target genes in *Gata2/Asxl1* double haploinsufficient HSCs. Yet, 16% (421) of all DEGs were unique to *Gata2/Asxl1* double haploinsufficient HSCs, indicating that combined *Gata2 and Asxl1* haploinsufficiency can in parallel generate a transcriptomic signature in HSCs distinct from that observed in either *Gata2^+/fl^; Vav-iCre^+^* HSCs or *Asxl1^+/fl^; Vav-iCre^+^* HSCs (**Supplementary Figure 6C**).

### *Gata2/Asxl1* double haploinsufficient HSCs display deregulated multi-lineage differentiation, membrane transport protein expression, DNA damage responses, inflammation and metabolic programmes

Using ICP analysis, we initially surveyed the entire transcriptomic signature of 2606 DEGs identified in *Gata2/Asxl1* double haploinsufficient HSCs, which consisted of DEGs that overlapped with *Gata2* and *Asxl1* single haploinsufficient HSCs. Consistent with hematopoietic differentiation defects observed in pre-leukemia (33, 41, 67, 68), transcriptional programmes specifying multi-lineage differentiation capacity were highly deregulated in *Gata2/Asxl1* double haploinsufficient HSCs (**Figure 6A**). Myriad solute carrier (SLC) membrane transport proteins were both up and down-regulated in *Gata2/Asxl1* double haploinsufficient HSCs, which likely reflects the paradoxical function of various SLCs to import nutrient substrates across cellular membranes in both normal and cancer settings (69) (**Figure 6A**). In *Gata2/Asxl1* double haploinsufficient HSCs a smaller selection of membrane ABC proteins, which, in contrast, act as export transporters to protect cells from undesirable metabolites and toxins, were upregulated (69, 70) (**Figure 6A**). In a similar vein to the transcriptional program in *Gata2* haploinsufficient HSC (**Figure 3** **and Supplementary Figure 3**), DNA damage and repair pathways (chromosomal organization genes, nucleotide excision repair, cell cycle checkpoints, G1/S regulation and cell cycle regulation by BTG) and inflammatory pathways (Death receptor signalling) were predominantly deregulated in *Gata2/Asxl1* double haploinsufficient HSCs (**Figure 6B**). Reflecting heightened cell cycling potential and impairment of DNA damage responses in *Gata2/Asxl1* double haploinsufficient HSCs, a proliferative transcriptomic signature was observed with up-regulation in pathways correlating with cell cycle regulation/cyclins and estrogen-mediated S-phase (**Figure 6B**). Metabolic pathways related to pro-oncogenic/AML activity were downregulated in *Gata2/Asxl1* double haploinsufficient HSCs. These included ketogenesis (where ketone bodies are produced in response to glucose deprivation), mevalonate pathway (which is involved in cholesterol metabolism), and glutathione detoxification (where glutathione acts a potent cellular antioxidant) (71–73) (**Figure 6B**). Therefore, *Gata2/Asxl1* double haploinsufficiency in HSCs causes wide-ranging transcriptional defects in DNA damage pathways, inflammation and metabolic pathways, which likely reflects genetic crosstalk between *Gata2* and *Asxl1* mediated biological pathways.

**Figure 6.**
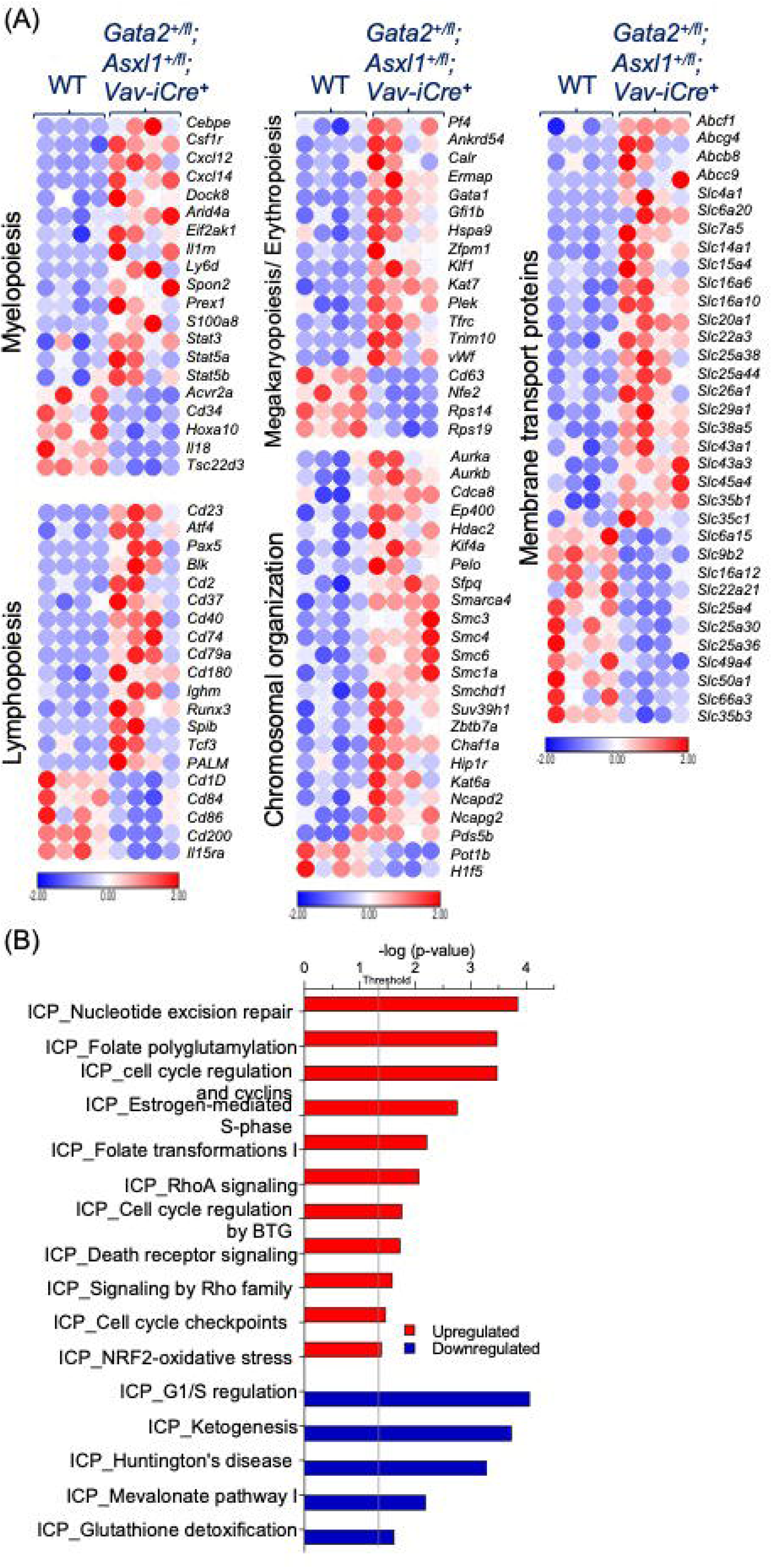
Transcriptional signature of *Gata2/Asxl1* double haploinsufficient HSCs reveals deregulated multi-lineage differentiation, membrane transport protein expression, DNA damage responses, inflammation, and metabolic programmes. RNA-Seq of purified HSCs (LSKCD150^+^CD48^-^) from *Gata2^+/fl^; Asxl1^+/fl^; Vav-iCre^-^*, *Asxl1^+/fl^; Vav-iCre^+^*, *Gata2^+/fl^; Vav-iCre^+^*, and *Gata2^+/fl^; Asxl1^+/fl^; Vav-iCre^+^*. n= 4 mice for each genotype. (A) Heatmaps of deregulated genes (DEseq2 computing-software, *P* <0.05 and FDR <0.05) in double haploinsufficient HSCs related to control HSCs. Analysis was performed using Morpheus online software showing a z-score scale. The red colour indicates high expression levels, and blue colour indicates low expression levels. (B) Canonical pathways of up-and down-regulated biological processes of dysregulated genes (DEseq2 computing-software, *P* <0.05 and FDR <0.05) in double haploinsufficient HSCs (n=4) compared with control HSCs (n=4) using IPA software. Data are presented as –log_10_ (p-value). The grey line indicates *P* value of 0.05. ICP indicates Ingenuity Canonical Pathway. Statistical analysis: Fischer’s Exact Test.

### Common GATA2 and ASXL1 target genes enforce an oncogenic and proliferation-biased transcriptional signature to *Gata2/Asxl1* double haploinsufficient HSCs

Next, we asked which shared *Gata2* and *Asxl1* biological pathways are involved in the 39 DEG core transcriptional signature found in *Gata2/Asxl1* double haploinsufficient HSCs, *Gata2* haploinsufficient HSCs, and *Asxl1* haploinsufficient HSCs. ICP analysis uncovered common biological pathways affected in each genotype were associated with DNA damage repair, hematopoietic differentiation to B-cell/myeloid lineages and deregulation of genes critical to development of hematopoietic neoplasms (**Figure 7A** and **B**). These data demonstrate that *Gata2/Asxl1* double haploinsufficient HSCs deregulate *Gata2* and *Asxl1* biological pathways essential for maintaining genomic and functional integrity of HSCs.

**Figure 7.**
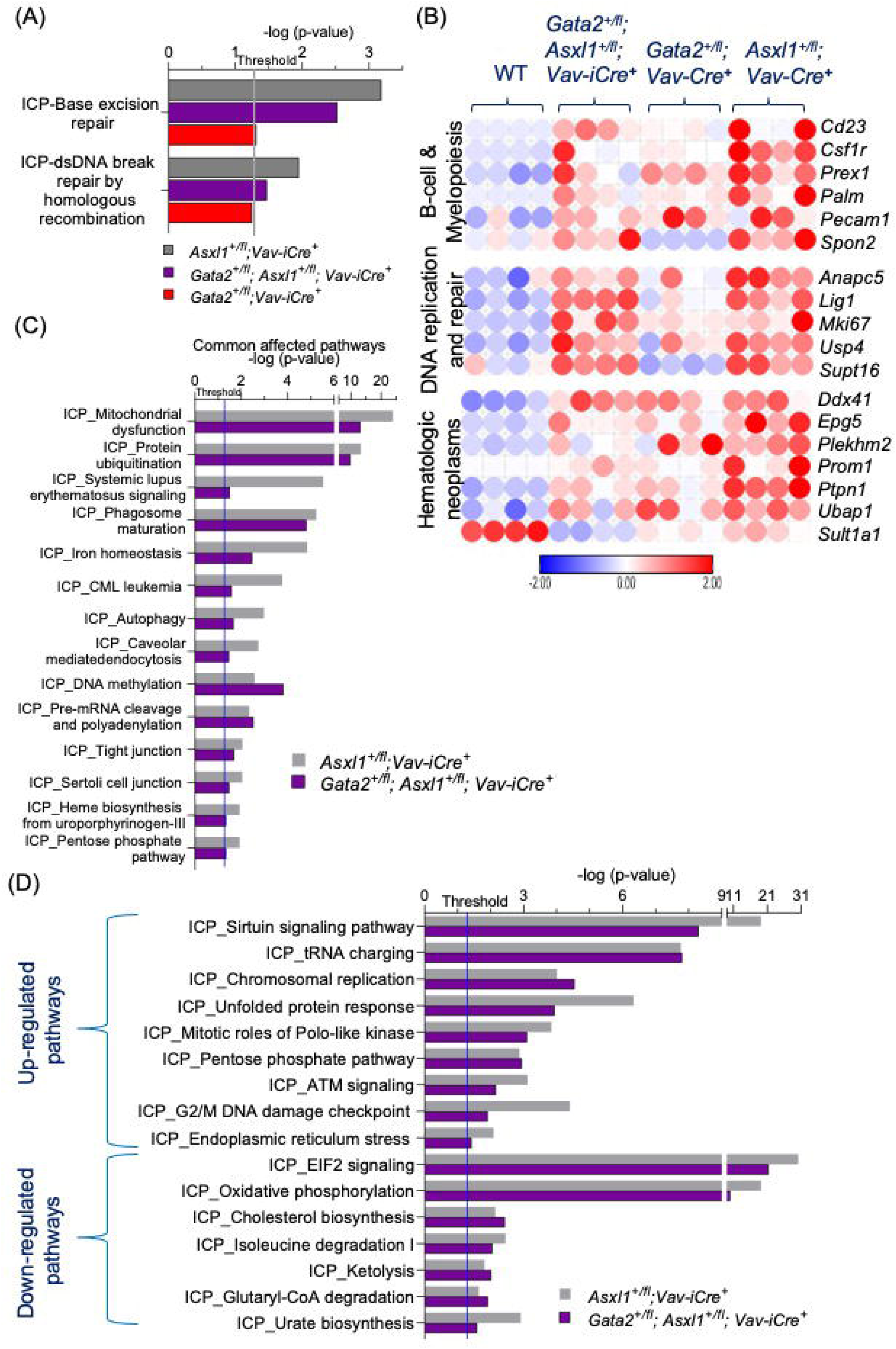
Common GATA2 and ASXL1 target genes embed an oncogenic and proliferation-biased transcriptional signature to *Gata2/Asxl1* double haploinsufficient HSCs. RNA-Seq of purified HSCs (LSKCD150^+^CD48^-^) from *Gata2^+/fl^; Asxl1^+/fl^; Vav-iCre^-^*, *Asxl1^+/fl^; Vav-iCre^+^*, *Gata2^+/fl^; Vav-iCre^+^*, and *Gata2^+/fl^; Asxl1^+/fl^; Vav-iCre^+^*. n= 4 mice for each genotype. (A) Biological pathway analysis of shared dysregulated pathways in *Gata2* haploinsufficient HSCs, *Asxl1* haploinsufficient HSCs, and double haploinsufficient HSCs. Presented data are – log_10_ (p-value), and the grey line represents the threshold at a p-value of 0.05. Fischer’s Exact Test. (B) Heatmaps of shared deregulated genes in *Gata2* haploinsufficient HSCs, *Asxl1* haploinsufficient HSCs, and double haploinsufficient HSCs. Heatmaps are drawn by Morpheus online software showing a z-score scale. The red colour indicates up-regulated genes; and blue colour, down-regulated genes. (C) IPA analysis of shared biological processes of dysregulated genes (DEseq2 computing-software, *P* <0.05 and FDR <0.05) in *Asxl1* haploinsufficient HSCs and double haploinsufficient HSCs compared with control HSCs. Presented data are –log_10_ (p-value). The blue line indicates a *P* value of 0.05. Fischer’s Exact Test. (D) Up- and down-regulated biological processes of shared dysregulated genes in *Asxl1* haploinsufficient HSCs, and double haploinsufficient HSCs using IPA software. Presented data are shown as –log_10_ (p-value), and the grey line indicates a *P* value of 0.05. ICP indicates Ingenuity Canonical Pathway. Fischer’s Exact Test.

ICP analysis revealed that certain biological pathways (e.g. DNA repair) were enhanced in *Gata2/Asxl1* double haploinsufficient HSCs beyond that found in single haploinsufficiency HSCs (**Figure 7A**). By complementary bioinformatic analysis using GSEA, we explored whether *Gata2* or *Asxl1* target genes/pathways were upregulated by *Gata2/Asxl1* double haploinsufficient HSCs. Target genes of MYC, a proto-oncogene involved in maintenance of HSC integrity and AML development (74), were highly enriched in double *Gata2/Asxl1* haploinsufficient HSCs compared to either *Gata2* or *Asxl1* haploinsufficient HSCs (**Supplementary Figure 7A and B**). In *Gata2/Asxl1* double haploinsufficient mice, G2M checkpoints, E2F target genes and mitotic spindle pathways, all of which are required to maintain genome instability (75), were upregulated compared to *Gata2* haploinsufficient HSCs (**Supplementary Figure 7B**). This finding is also consistent with the observed increased HSCs cycling status in these mice (**Figure 5**). Furthermore, spliceosome and adipogenesis pathways, both of which are deregulated in the context of MDS/AML (76, 77), were enriched in *Gata2/Asxl1* double haploinsufficient HSCs compared with *Gata2* haploinsufficient HSCs (**Supplementary Figure 7B**).

As approximately 50% of DEGs were shared between *Gata2/Asxl1* double haploinsufficient HSCs and *Asxl1^+/fl^; Vav-iCre^+^* HSCs, we also explored the commonality in biological pathways between these two genotypes. By ICP analysis, common affected pathways in *Gata2/Asxl1* double haploinsufficient HSCs and *Asxl1^+/fl^; Vav-iCre^+^*HSCs broadly encompassed metabolism (mitochondrial dysfunction), protein ubiquitination, immune cell homeostasis (phagosome maturation), cellular stress responses (autophagy), DNA methylation and RNA-processing pathways (**Figure 7C**). By segregating these common affected pathways into specific up-regulated and downregulated pathways in ICP analysis, we found G2M damage checkpoints, ATM signalling, ER stress/unfolded protein response (UPR) and pentose phosphate pathway were amongst some key pathways that were up-regulated in *Gata2/Asxl1* double haploinsufficient HSCs and *Asxl1^+/fl^; Vav-iCre^+^* HSCs (**Figure 7D**). Likewise, essential metabolic pathways such as cholesterol biosynthesis, urate biosynthesis, isoleucine degradation, and ketolysis were down-regulated in *Gata2/Asxl1* double haploinsufficient HSCs and *Asxl1^+/fl^; Vav-iCre^+^* HSCs (**Figure 7D**). Notably, the common feature of affected, upregulated and downregulated pathways in these two genotypes was their role in MDS/AML pathogenesis (75, 78–80). Overall, this analysis suggests that common GATA2 and ASXL1 target genes/pathways in *Gata2/Asxl1* double haploinsufficient HSCs co-operate to fortify an oncogenic and proliferation-biased transcriptomic program.

### *Gata2/Asxl1* double haploinsufficient HSCs formulate a unique impaired DNA damage and inflammatory response, disrupted amino acid metabolism and a pre-leukemic transcriptional network

Having established common GATA2 and ASXL target genes/pathways in *Gata2/Asxl1* double haploinsufficient HSCs, we conducted separate pathway analysis on the unique 421 DEGs identified in *Gata2/Asxl1* double haploinsufficient HSCs. Analysis of these DEGs in ICP and GSEA databases unveiled biologic pathways recurrently linked to the HSC transcriptional signature of all genotypes, namely DNA damage responses/proliferation (GADD45 signalling, MYC target genes, apoptosis and mitotic spindle) and inflammatory responses (neutrophil degranulation, IFN-D response, and Tnfα signalling via NF-κB) (**Figure 8A**). By utilizing both *Gata2/Asxl1* dependent and independent mechanisms, impaired DNA damage and inflammatory responses therefore appear to be critical to the formulation of the *Gata2/Asxl1* double haploinsufficient HSC transcriptional network. Amino-acid metabolism pathways were also deregulated in the unique 421 DEGs identified in *Gata2/Asxl1* double haploinsufficient HSCs (**Figure 8A**). Two amino-acid metabolic pathways affected in *Gata2/Asxl1* double haploinsufficient HSCs are particularly noteworthy for their specific role in AML. First, biosynthesis of cysteine, a non-essential amino acid required for protein synthesis and regulation of oxidative stress, is essential for AML cell survival (81). Second, methionine degradation related pathways include the conversion of methionine to S-adenosyl-methionine, an intermediate that is involved in DNA methylation, which is an affected pathway in both *Gata2/Asxl1* double haploinsufficient HSCs and *Asxl1* single haploinsufficient HSCs (**Figure 8A**) (82, 83). Furthermore, S-adenosyl-methionine is overexpressed in poor prognosis AML (83). Finally, we also found evidence for *Gata2/Asxl1* double haploinsufficient HSCs specific DEGs contributing to a robust gene signature relating more specifically to MDS. This signature consisted of deregulated *Prmt5*, *Hmgcs1*, *Lamp2*, *Gdf15* and *Mlf1* expression (84–86) as well as aberrant Wnt signalling (87) and mRNA spliceosome pathways (pre-mRNA splicing) (88) (**Figure 8A** and **B**). These data indicate that *Gata2/Asxl1* double haploinsufficient HSCs express a unique pre-leukemic transcriptional signature, which may be germane to the progression of clinical *GATA2* haploinsufficiency syndromes.

**Figure 8.**
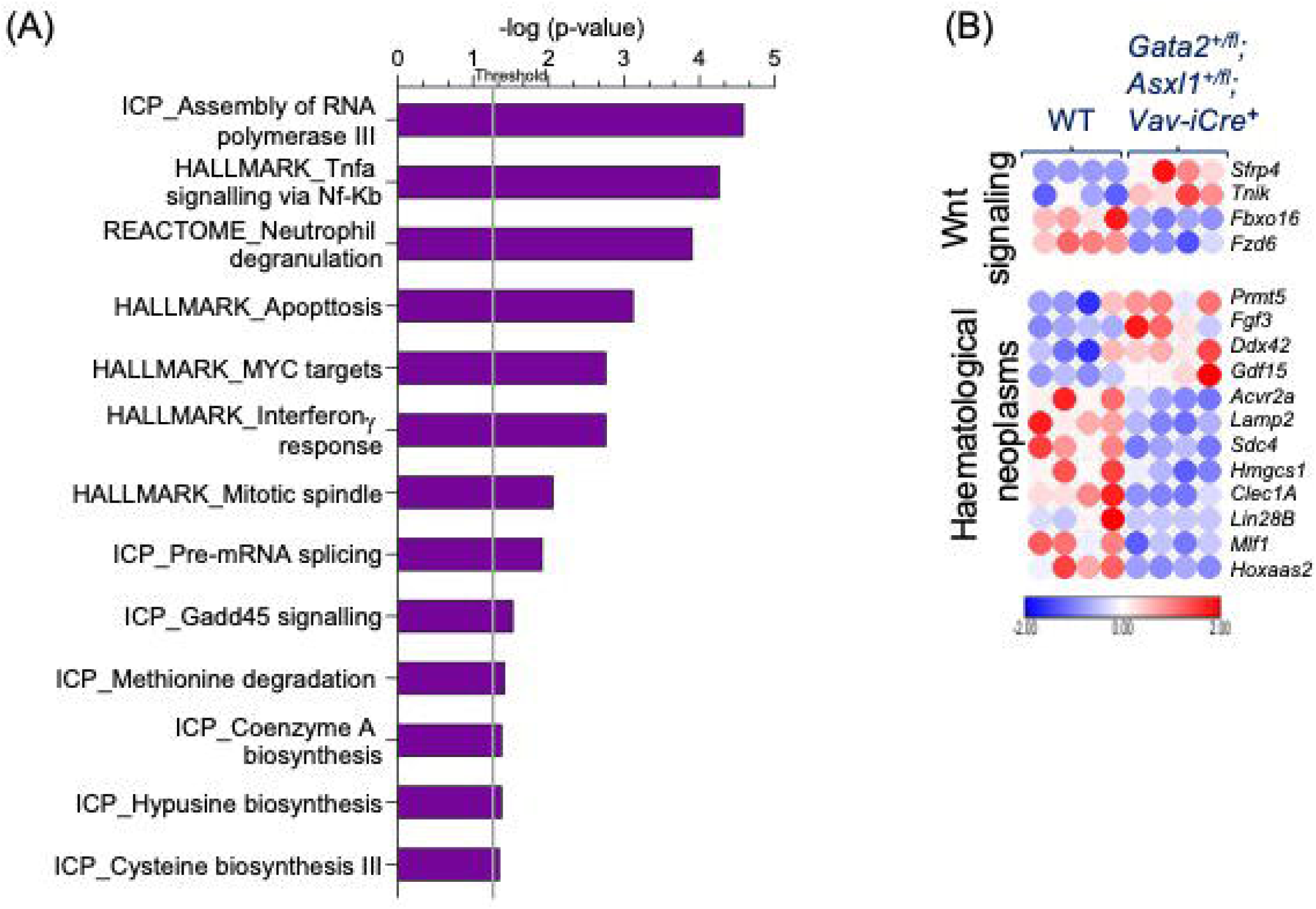
Unique transcriptional signature of double *Gata2*/*Asxl1* haploinsufficient HSCs formulates an impaired DNA damage, inflammatory response, disrupted amino acid metabolism and a MDS transcriptional network. RNA-Seq analysis of sorted HSCs (LSKCD150^+^CD48^-^) from *Gata2^+/fl^; Asxl1^+/fl^; Vav-iCre^-^*, *Asxl1^+/fl^; Vav-iCre^+^*, *Gata2^+/fl^; Vav-iCre^+^*, and *Gata2^+/fl^; Asxl1^+/fl^; Vav-iCre^+^*. n= 4 mice for each genotype. (A and B) IPA, HALLMARK, and REACTOME analyses of biological pathways (A) and heatmaps (B) of 421 unique dysregulated genes (*P* <0.05 and FDR <0.05) in double haploinsufficient HSCs related to control HSCs. Data are presented as –log_10_ (p-value). The grey line signifies the *P* value of 0.05 level. Heatmaps show a z-score scale. Statistical analysis: Fischer’s Exact Test.

## Discussion

Predating the discovery of familial *GATA2* haploinsufficiency syndromes, where patients develop primary immunodeficiency syndromes that transform to MDS/AML, mouse models of germline *Gata2* haploinsufficiency demonstrated underlying HSC/HSPC functional defects (9, 36). Using highly sophisticated genetic mouse models, further work since then has demonstrated that *Gata2* cis-regulatory elements, which are mutated in *GATA2* deficient patients, are essential for HSC genesis in the embryo and necessary for HSPC regeneration in response to genotoxic insults (24, 89). Yet our understanding of the functional and transcriptional program mediated by *GATA2* haploinsufficiency in HSCs/HSPCs remains ill-defined. Furthermore, the functional impact of secondary genetic mutations acquired by *GATA2* deficient HSCs/HPCs during disease progression and, indeed, their involvement in shifting the disease from immunodeficiency to the pre-leukemic state is poorly characterised. We addressed these issues using a conditional genetic knockout of *Gata2* mediated by *Vav-Cre* and found that young adult *Gata2* haploinsufficient mice display HSC functional defects in self-renewal and multilineage differentiation capability. *Gata2* haploinsufficiency leads to defects in HSPC lineage specification during B-cell maturation, early erythroid commitment, maturation of platelets and inflammatory cell generation, all of which is underpinned by deregulated DNA damage responses and inflammatory signalling emanating from *Gata2* haploinsufficient HSCs. Analysis of compound young adult *Gata2/Asxl1* haploinsufficient mice demonstrated that *Asxl1*, a secondary mutation driving MDS/AML in about 30% of familial *GATA2* haploinsufficiency syndromes (33, 34), interacts genetically with *Gata2* to potentiate deregulated DNA damage and inflammatory signalling in HSCs that was initiated in *Gata2* haploinsufficient mice, and, through both unique and GATA2/ASXL1 dependent mechanisms, develop a robust pre-leukemic transcriptional program. These data present a complex portrait of how *Gata2* haploinsufficiency itself initially drives deregulation of HSC genome stability and explain how this could, on the acquisition of secondary mutations like *ASXL1*, lead to an increased susceptibility to transform *GATA2* haploinsufficiency syndromes from immunodeficiency into pre-leukemia.

*GATA2* haploinsufficient patients display both myeloid and lymphoid lineage specific defects and previous *Gata2* haploinsufficient mouse models have been partially successful in modelling these defects linked to myeloid lineage commitment. In this report, in addition to the expected reduction in HSCs and progenitors related to the myeloid lineage in *Gata2* haploinsufficient mice (9, 11), we identified selective B-cell development blocks in BM and spleen, which was preceded by attenuation in lymphoid primed progenitor (LMPP/HPC1) numbers and deregulation of the B-cell transcriptional program that begins in HSCs. While we did not observe defects in mature B-cells in *Gata2* haploinsufficient mice, this data provides the basis for understanding the differentiation stage specific dependency for GATA2 in B-cell maturation in *GATA2* haploinsufficient patients, where the incipient stages of B-cell differentiation from HSCs/multi-potent progenitors are known to be disrupted (37–39).

We further identified megakaryocytic lineage specific alterations in *Vav-Cre* mediated *Gata2* haploinsufficiency, which has a provenance within MPPs. MPP2, a megakaryocyte/erythroid-biased MPP compartment, was expanded whereas downstream failure of megakaryocyte/erythroid committed progenitors to mature into megakaryocytes likely caused the reduction in platelets observed in *Gata2* haploinsufficient mice. Despite defects in early erythroid commitment, evidenced by PreMeg/E and Pre-CFU-E reduction in *Gata2* haploinsufficient mice, later erythropoiesis, as measured by CFU-E/Pro-E, was broadly unimpacted by Gata2 haploinsufficiency, likely due to compensation from other indispensable erythroid factors including GATA family member, *Gata1* (90). Overall, these data support studies implicating a pivotal role for *Gata2* in megakaryocyte differentiation (13, 91). Since HSCs can directly generate platelets independently of MPPs (92), we cannot exclude the possibility of a parallel reduction in platelets caused exclusively by HSC reduction observed in *Vav-Cre* mediated *Gata2* haploinsufficiency. Given that megakaryocytes regulate HSCs quiescence by secreting important factors as CXCL4, TPO, and TGFβ1 (93–95), enhanced HSC quiescence observed in *Gata2* haploinsufficient mice appears to be independent of megakaryocytes, as the observed reduction in megakaryocytes would instead be predicted to drive HSC proliferation (91). Notably, our model of *Gata2* haploinsufficiency reflects crucial features of deregulated megakaryocyte biology and early erythroid commitment observed in *GATA2* deficiency patients including skewed HSCs differentiation toward megakaryocyte/erythrocyte progenitors in MDS driven familial *GATA2* haploinsufficiency (96), frequently observed BM erythroid /megakaryocytic features of dysplasia in *GATA2* haploinsufficiency patients and peripheral thrombocytopenia, seen in about 20% of patients harbouring *GATA2* mutations that have evolved to MDS/AML (6, 13, 39, 97).

Functional analysis of *Gata2* haploinsufficient HSCs in transplantation experiments revealed defects in self-renewal and multilineage differentiation capability. Underlying these HSC functional defects was impaired DNA damage responses (DDR). Of relevance to *GATA2* haploinsufficiency, defective DDR activities can lead to BM failure, immunodeficiency, and hematological disorders (98–100). Given that damaged DNA in quiescent HSCs is corrected by the error prone nonhomologous end-joining (NHEJ) repair pathway that potentially enhances mutagenesis (101, 102), it is possible that enhanced HSC quiescence observed in *Gata2* haploinsufficient mice (9, 40) makes them more susceptible to acquisition of secondary mutations that drive malignant transformation (101). Since the G0/G1 checkpoint is the major restriction checkpoint before entering mitosis (103), in the setting of *Gata2* haploinsufficiency NHEJ could potentially expand the pool of quiescent HSCs with genomic instability, and, if those HSCs are not completely eliminated by apoptosis, on re-entry into cycle they can export genomic instability to both proliferating and differentiating HSPC pools. Together with deregulation of both high-fidelity DDR via HR, which in contrast occurs exclusively in proliferating cells (102), and base-excision repair, our data suggests genome instability builds across both quiescent and proliferating HSC compartments in *Gata2* haploinsufficiency. Up-regulation of HSC apoptosis observed in *Gata2* haploinsufficient HSCs (9, 40) may be a safeguard to preserve HSC genomic integrity, yet it also further supports the notion of inefficient DDR mechanisms operating within HSCs (98–100). Future investigations will be required to identify the critical DNA damage molecular pathways deregulated in NHEJ and HR in *Gata2* haploinsufficient HSCs and how this contributes to differentiation specific impacts in HSPCs and their downstream progeny.

HSCs from *Gata2^+/fl^; Vav-iCre^+^* mice demonstrated deregulated inflammatory signalling, typified by up-regulation of IFN-γ signalling that may explain the observed HSC self-renewal defect (59, 60). It is also of interest to contemplate whether enhanced inflammatory signalling in *Gata2^+/fl^; Vav-iCre^+^*HSC is driven by the deregulated DNA damage response or *vice versa*. In the former scenario, induced quiescence and apoptosis in *Gata2^+/fl^; Vav-iCre^+^* HSCs in response to DNA damage stimuli are known to elicit inflammatory cytokine secretion (59, 60, 98). Alternatively, the chronic inflammatory tone evident in HSCs may initiate DNA damage genomic instability via the production of reactive oxygen species (ROS) (59, 60, 98). In whatever order deregulated DNA damage and inflammatory signalling pathways happens in *Gata2^+/fl^; Vav-iCre^+^* HSCs, a perpetual feedback loop and crosstalk may be maintained between these pathways, thereby promoting mutagenesis (59, 60, 98). Further studies to elucidate the mechanistic relationship between genome stability and inflammation in *Gata2^+/fl^; Vav-iCre^+^* mice will be of interest.

Surprisingly, the enhanced inflammatory tone in HSCs shifts to the production of anti-inflammatory cells in the BM/spleen and down-regulation of pro-inflammatory cells in the spleen of *Gata2^+/fl^; Vav-iCre^+^* mice. Notably, however, altered production of macrophage inflammatory subsets is likely consistent with monocytopenia in *GATA2* haploinsufficiency patients and impaired inflammatory responses to pathogenic insults observed in *Gata2* haploinsufficient mice and patients alike (26, 27, 32, 63). Why this occurs is unclear, but there are several possible explanations. Anti-inflammatory cell generation or down-regulation of pro-inflammatory cells during HSC differentiation may be a compensatory mechanism to correct for enhanced inflammatory signalling from *Gata2* haploinsufficient HSCs. Second, the anti-inflammatory cell feedback may operate to preserve the growth and expansion of some abnormal HSC clones, whereas overall growth of *Gata2* haploinsufficient HSCs via self-renewal is curtailed (104). Finally, heightened inflammatory signalling in *Gata2* haploinsufficient HSCs may simply cause disordered differentiation to inflammatory cell lineages in the BM and spleen (59, 60).

We conducted analysis of *Gata2/Asxl1* double haploinsufficient mice and shRNAi *Asxl1* knockdown experiments in *Gata2^+/fl^; Vav-iCre^+^* HSCs in order to understand how secondary mutations capitalize on genomic instability and deregulate inflammatory signalling in *Gata2^+/fl^; Vav-iCre^+^* HSCs to facilitate the evolution of *GATA2* haploinsufficiency to MDS/AML. *Gata2/Asxl1* double haploinsufficient mice largely phenocopied the HSPC characteristics of *Gata2^+/fl^; Vav-iCre^+^*HSPCs and transcriptomic signatures that overlapped considerably with their single heterozygote counterparts, including DNA damage response and inflammatory signalling. This suggests that *Gata2* and *Asxl1* cooperate to reenforce genomic instability via DNA damage responses and inflammatory signalling. However, in contrast to their single heterozygote counterparts, *Gata2/Asxl1* double haploinsufficient mice HSCs were more proliferative and engrafted poorly in transplantation experiments. Hyper-proliferation of *Gata2^+/fl^; Vav-iCre^+^* HSCs was consistent with progression through cell cycle, supported by transcriptomic profiling indicating increased deregulation of key later stage cell cycle regulators (e.g. G1/S, G2M checkpoints, DNA replication initiators and mitosis related pathways) linked to an increased oncogenic potential. For example, MYC target genes were enhanced in *Gata2/Asxl1* double haploinsufficient HSCs and are implicated in cell cycle regulation, yet persistent expression of MYC target genes also positively correlates with tumorigenesis (105, 106). Thus, our data point to HSC proliferation enabling the development of a pre-leukemia transcriptomic signature in young *Gata2/Asxl1* double haploinsufficient mice, an idea that is further supported by the unique MDS transcriptional signature observed in young *Gata2/Asxl1* double haploinsufficient mice and functional assays showing a differentiation block to committed progenitors in CFCs from shRNA *Asxl1* knockdown experiments in *Gata2^+/fl^; Vav-iCre^+^* HSCs and following transplantation of HSCs from *Gata2/Asxl1* double haploinsufficient mice. Future work will be focused on evaluating hematopoietic potential and disease progression in aged *Gata2/Asxl1* double haploinsufficient mice.

*GATA2* haploinsufficient patients with somatic *ASXL1* mutations have a rapid onset of MDS/AML and poor prognosis (33, 34, 41). Our study offers insight into the mechanisms which may operate to facilitate progression of clinical *GATA2* haploinsufficiency syndromes from immunodeficiency to hematologic malignancy and provide a framework to understand the basis for poorer prognosis in these patients. A case in point is our finding that ABC export proteins were upregulated in *Gata2/Asxl1* double haploinsufficient HSCs, providing a plausible basis for chemoresistance in these patients (69, 70) and an opportunity for pharmacologic intervention to eradicate pre-leukemic/leukemic stem cells (69, 70). Thus, the wealth of transcriptomic data presented in our analysis of young *Gata2/Asxl1* double haploinsufficient HSCs should also inform early therapeutic interventions to overcome the challenge of poor prognosis in these patients.

## Conflicts of Interest Disclosure

The authors declare no conflicts of interest.

## Author contribution

A. Abdelfattah designed and performed experiments, analyzed and interpreted data (including RNA-seq), prepared the figures, and contributed to writing the manuscript., A.H., L-aT, J.B.M-G and A. Almotiri performed experiments, analyzed data, and provided critical input during manuscript preparation, H. Alqahtani, H.L., S.T., M.S., R.A., H. Alzahrani and A.C. performed experiments and analyzed data., A.G. and P.G. conducted initial RNA-seq anlaysis, A.T. contributed to experimental design and analysis and provided critical review of the manuscript, A.S.B. and K.R.K. contributed significantly to experimental design, data analysis/interpretation and contributed to manuscript preparation/critical review of manuscript and NPR conceived and supervised the project, designed experiments, analyzed and interpreted the data, and wrote the manuscript.

## Acknowledgements

The Rodrigues laboratory at Cardiff University, Cardiff, UK is supported by the Saudi Arabia Cultural Bureau, Blood Cancer UK, Leukaemia Cancer Society, and Leukaemia and Myeloma Research UK. We would like to thank the Deanship of Scientific Research at Shaqra University, Shaqra, Saudi Arabia for supporting the work of A. Almotiri and The Hashemite University, Zarqa, Jordan for supporting the work of A. Abdelfattah.

## Supplementary Figure Legends

**Supplementary Figure 1, related to Figure 1. *Gata2* haploinsufficiency impairs megakaryopoiesis and committed myeloid progenitors.** (A) Representative agarose gel electrophoresis image shows efficient excision of the floxed *Gata2* allele in BM cells. fl indicates floxed allele; +, wildtype allele; and bp, base pair. (B) qRT-PCR showing relative expression of *Gata2* mRNA compared with *Hprt* mRNA in BM and purified LSK cells from control and *Gata2^+/fl^; Vav-iCre^+^* mice using the comparative 2^-ΔΔCT^ method. n= 3 mice for each genotype. (C) Analysis of total cellularity numbers in BM (from two femurs and tibias, cells/30mL), spleen (cells/7mL), and thymus (cells/4mL) in control (n=6) and *Gata2^+/fl^; Vav-iCre^+^* (n=7) mice from three independent experiments. (D) Frequency of differentiated myeloid (Mac1 and Gr1) and lymphoid (B220, CD4, and CD8) cells in the PB from control and *Gata2^+/fl^; Vav-iCre^+^* mice from three independent experiments. n= 6 for each genotype. (E) Frequency of different T-lymphocyte populations in the thymus of CD3, CD4, CD8, DN (CD4^−^CD8^−^), and DP (CD4^+^CD8^+^) from control (n = 6) and *Gata2^+/fl^; Vav-iCre^−^* (n = 7) mice. Data were collected from three independent experiments. (F and G) Percentage of cells in the BM (F) and spleen (G) of myeloid (Mac1^+^ Gr1^+^), erythroid (proerythroblasts, CD71 Ter119 ^−/low^; early-erythroblasts, CD71 Ter119^−/low^; and late erythroblasts, CD71^−^ Ter119^+^); B-lymphoid (B220^+^), and T-lymphoid (CD4^+^ and CD8^+^) cells from control (n=6) and *Gata2^+/fl^; Vav-iCre^−^* (n=7) mice from three independent experiments. (H and I) Representative immunophenotypic analysis (H) and absolute numbers (I) of BM common lymphoid progenitors (CLP, Lin^−^c-kit*lo* Sca1^lo^ CD127^+^) from control (n = 6) and *Gata2^+/fl^; Vav-iCre^−^* (n = 7) mice from three independent experiments. Data are presented as mean ± SEM. Statistical analysis: Mann-Whitney U test.

**Supplementary Figure 2, related to Figure 2. *Gata2* haploinsufficiency affects the development of primitive hematopoietic compartments and impairs multi-lineage reconstitution and self-renewal capacity of adult HSCs after transplantation.** (A and B) Percentage of total CD45.2 cells and donor contribution in the BM (A) and spleen (B) for myeloid-cells (Mac1^+^Gr1^+^), erythroid-cells (Ter119^+^) T-cell (CD4^+^ and CD8^+^), and B-cells (B220^+^) at week 16 after primary HSCs transplantation from control (n=8) and *Gata2^+/fl^; Vav-iCre^+^* (n=8) derived cells from 3 independent experiments. (C) Proportion of total CD45.2 cells in the thymus and donor contribution to different T-lymphocyte subsets of CD3, CD4, CD8, DN (CD4^−^CD8^−^), and DP (CD4^+^CD8^+^) at week 16 post primary HSCs transplantation from control (n=6) and *Gata2^+/fl^; Vav-iCre^+^* (n=6) derived cells from 3 separate experiments. (D) Frequency of CD45.2 cells in the BM at week 20 after secondary HSCs transplantation from primary recipients of control (n= 7) *Gata2^+/fl^; Vav-Cre^+^* (n= 7) derived cells from 3 independent experiments. Presented data are mean ± SEM. Statistical analysis: Mann-Whitney U test *, *P* < 0.05; **, *P* < 0.01; ***, *P* < 0.001.

**Supplementary Figure 3, related to Figure 3. *Gata2* haploinsufficient HSCs display deregulation of transcriptional signatures associated with the DNA damage repair and proinflammatory signalling.** (A) Venn diagrams illustrate significant numbers of differentially dysregulated genes in purified HSCs from *Gata2^+/fl^; Vav-iCre^+^* (n= 4) mice as compared to control HSCs (n= 4) using DEseq2 computing-software with p-values and FDR-values of less than 0.05 level. (B) Biological and molecular processes for differentially expressed genes in *Gata2* haploinsufficient HSCs relative to control HSCs. Data are presented as –log_10_ (p-value), and the grey line indicates a p-value of 0.05, using Fischer’s Exact Test. (C) Heatmaps of the differentially deregulated genes in *Gata2* haploinsufficient HSCs compared with control HSCs. Heatmaps are drawn by Morpheus online software and show a z-score scale. The red colour indicates up-regulated genes, while the blue colour indicates down-regulated genes. (D and E) Representative immunophenotypic analysis (D) and percentages (J) of B lymphocyte precursors in the BM from control (n= 9) and *Gata2 ^+/fl^; Vav-iCre^+^* (n= 9) mice. Early B cell precursors (CD43+IgM-) differentiate into pre-pro B cells (BP1-CD19-), pro-B cells (BP1-CD19+), and early pre-B cells (BP1+CD19+). Late pre-B cells (CD43-IgM-) and immature B cells (CD43-IgM+) are distinguished depending on CD19 expression. and. Data are presented as mean ± SEM. Statistical analysis: Mann-Whitney U test. (F) GSEA analysis displaying enrichment of up-regulated genes in *Gata2* haploinsufficient HSCs related to control HSCs. NES (normalised enrichment score), *P*, and FDR values are shown. (G) Percentages of phagocytic and pro-inflammatory macrophages (Mac1^+^ Ly6G^−^ CD115^+^ Ly6C^high^), pro-inflammatory macrophages (Mac1^+^ Ly6G^−^ CD115^+^ Ly6C^low^) and anti-inflammatory macrophages (Mac1^+^ Ly6G^−^ CD115^+^ Ly6C^low^) in spleen from control and *Gata2^+/fl^; Vav-iCre^−^* mice from two independent experiments. Data are presented as mean ± SEM. Statistical analysis: Mann-Whitney U test.

**Supplementary Figure 4, related to Figure 5. *Gata2*/*Asxl1* double haploinsufficient mice demonstrated increased HSCs cycling, decreased HSCs survival, and ultimately exhausted HSC pool size after HSCs transplantation.** (A) Representative genomic DNA analysis indicates efficient excision of the floxed allele of *Gata2* and *Asxl1* in BM cells from *Gata2^+/fl^; Asxl1^+/fl^; Vav-iCre^+^* mice. fl indicates floxed allele; +, wildtype allele; and bp, base pair. (B) *Asxl1* mRNA expression in total BM cells from control (n=5) and *Asxl1^+/fl^; Vav-iCre^+^* (n=3) mice relative to *Hprt* mRNA. Mann-Whitney U test. (C) *Gata2* and *Asxl1* mRNA expression levels in BM c-kit^+^ cells from control (n=3) and *Gata2^+/fl^; Asxl1^+/fl^; Vav-iCre^+^* (n=2) littermates relative to *Hprt* mRNA. Mann-Whitney U test. (D) Absolute values of PB CBC indices (WBCs, RBCs, haemoglobin level, and platelets) from control (n=6), *Asxl1^+/fl^; Vav-iCre^+^* (n=7), *Gata2^+/fl^; Vav-iCre^+^* (n=6), and *Gata2^+/fl^; Asxl1^+/fl^; Vav-iCre^+^* (n=7) mice from two independent experiments. One-Way ANOVA with Tukey’s multiple comparisons test. (E) Frequency of myeloid (Mac1^+^Gr1^+^, Mac1^+^Gr1^-^) and lymphoid (B220^+^, CD4^+^, and CD8^+^) cells in the PB of control (n=10), *Asxl1^+/fl^; Vav-iCre^+^* (n=10), *Gata2^+/fl^; Vav-iCre^+^* (n=10), and *Gata2^+/fl^; Asxl1^+/fl^; Vav-iCre^+^* (n=10) mice from three independent experiments. One-Way ANOVA with Tukey’s multiple comparisons test. (F) Total cellularity numbers in BM (from two femurs and tibias, cells/30mL) and spleen (cells/7mL) from control, (n=8), *Asxl1^+/fl^; Vav-iCre^+^* (n=8), *Gata2^+/fl^; Vav-iCre^+^* (n=8), and *Gata2^+/fl^; Asxl1^+/fl^; Vav-iCre^+^* (n=8) mice from three independent experiments. One-Way ANOVA with Tukey’s multiple comparisons test. (G and H) Proportions of myeloid (Mac1^+^Gr1^+^), erythroid (Ter119^+^) and lymphoid (B220^+^, CD4^+^, and CD8^+^) cells in the BM (G) and spleen (H) from control (n=8), *Asxl1^+/fl^; Vav-iCre^+^* (n=8), *Gata2^+/fl^; Vav-iCre^+^* (n=8), and *Gata2^+/fl^; Asxl1^+/fl^; Vav-iCre^+^* (n=8) mice from three independent experiments. Data are presented as mean ± SEM. Statistical analysis: One-Way ANOVA with Tukey’s multiple comparisons test.

**Supplementary Figure 5, related to Figure 5. *Gata2*/*Asxl1* double haploinsufficient mice demonstrated increased HSCs cycling, decreased HSCs survival, and ultimately exhausted HSC pool size after HSCs transplantation.** (A) Follow-up analysis of total CD45.2^+^ cells in PB at different time points after HSCs transplantation from control (n=7), *Asxl1^+/fl^; Vav-iCre^+^* (n=6), *Gata2^+/fl^; Vav-iCre^+^* (n=6), and *Gata2^+/fl^; Asxl1^+/fl^; Vav-iCre^+^* (n=6) donor derived-cells from two independent experiments. (B) Proportion of CD45.2 donor cells in PB at week 16 post HSCs transplantation for myeloid (Mac1^+^Gr1^+^, Mac1^+^Gr1^-^) and lymphoid (B220^+^, CD4^+^, and CD8^+^) cells from control (n=7), *Asxl1^+/fl^; Vav-iCre^+^* (n=6), *Gata2^+/fl^; Vav-iCre^+^* (n=6), and *Gata2^+/fl^; Asxl1^+/fl^; Vav-iCre^+^* (n=6) donor derived-cells from two independent experiments. Data are presented as mean ± SEM. Statistical analysis: One-Way ANOVA with Tukey’s multiple comparisons test *, P < 0.05; **, P < 0.01; ***, P < 0.001; ****, P < 0.0001.

**Supplementary Figure 6. Transcriptional signature of *Asxl1* haploinsufficient HSCs reveals deregulated biological processes involved in hematological malignancies.** RNA-Seq of purified HSCs (LSKCD150^+^CD48^-^) from *Gata2^+/fl^; Asxl1^+/fl^; Vav-iCre^-^*, *Asxl1^+/fl^; Vav-iCre^+^*, *Gata2^+/fl^; Vav-iCre^+^*, and *Gata2^+/fl^; Asxl1^+/fl^; Vav-iCre^+^*. n= 4 mice for each genotype. (A and B) Venn diagrams of significantly up- and down-regulated genes in double haploinsufficient HSCs (A) and *Asxl1* haploinsufficient HSCs (B) compared to control HSCs. The identification of expressed genes was achieved using DEseq2 computing-software based on p-values and FDR-values of <0.05 level. (C) Numbers and overlap analysis of DEseq2 in *Gata2* haploinsufficient HSCs, *Asxl1* haploinsufficient HSCs, and double haploinsufficient HSCs using Fischer’s Exact Test. (D) RNA-Seq analysis of *Asxl1* haploinsufficient HSCs. IPA analysis of up-regulated and down-regulated biological pathways of dysregulated genes (DEseq2 computing-software, *P* <0.05 and FDR <0.05) in *Asxl1* haploinsufficient HSCs (n=4) compared with control HSCs (n=4). Data are shown as –log_10_ (p-value). The threshold line in grey shows a *P* value of 0.05. ICP indicates Ingenuity Canonical Pathway. Statistical analysis: Fischer’s Exact Test.

**Supplementary Figure 7, related to Figure 7. Common GATA2 and ASXL1 target genes embed an oncogenic and proliferation-biased transcriptional signature to *Gata2/Asxl1* double haploinsufficient HSCs.** RNA-Seq analysis of sorted HSCs (LSKCD150^+^CD48^-^) from *Gata2^+/fl^; Asxl1^+/fl^; Vav-iCre^-^*, *Asxl1^+/fl^; Vav-iCre^+^*, *Gata2^+/fl^; Vav-iCre^+^*, and *Gata2^+/fl^; Asxl1^+/fl^; Vav-iCre^+^*. n= 4 mice for each genotype. (A and B) GSEA analysis of genes enriched in double haploinsufficient HSCs compared with *Asxl1* haploinsufficient HSCs (A) and *Gata2* haploinsufficient HSCs (B) with *p* <0.05 and FDR <0.05. NES indicates normalised enrichment score.

## Notes

### Competing Interest Statement

The authors have declared no competing interest.

## References

1. Keohane EM, Smith L, and Walenga JM. Rodak’s hematology: clinical principles and applications. Elsevier Health Sciences; 2015.

2. Rieger MA, and Schroeder T. Hematopoiesis. Cold Spring Harb Perspect Biol. 2012;4(12):a008250.

3. Khoury JD, Solary E, Abla O, Akkari Y, Alaggio R, Apperley JF, et al. The 5th edition of the World Health Organization classification of haematolymphoid tumours: myeloid and histiocytic/dendritic neoplasms. Leukemia. 2022;36(7):1703–19.

4. Merika M, and Orkin SH. DNA-binding specificity of GATA family transcription factors. Molecular and cellular biology. 1993;13(7):3999–4010.

5. Tsai S-F, Martin DI, Zon LI, D’Andrea AD, Wong GG, and Orkin SH. Cloning of cDNA for the major DNA-binding protein of the erythroid lineage through expression in mammalian cells. Nature. 1989;339(6224):446–51.

6. Gao X, Johnson KD, Chang Y-I, Boyer ME, Dewey CN, Zhang J, et al. Gata2 cis-element is required for hematopoietic stem cell generation in the mammalian embryo. J Exp Med. 2013;210(13):2833–42.

7. de Pater E, Kaimakis P, Vink CS, Yokomizo T, Yamada-Inagawa T, van der Linden R, et al. Gata2 is required for HSC generation and survival. J Exp Med. 2013;210(13):2843–50.

8. Menendez-Gonzalez JB, Vukovic M, Abdelfattah A, Saleh L, Almotiri A, Thomas L-a, et al. Gata2 as a crucial regulator of stem cells in adult hematopoiesis and acute myeloid leukemia. Stem cell reports. 2019;13(2):291–306.

9. Rodrigues NP, Janzen V, Forkert R, Dombkowski DM, Boyd AS, Orkin SH, et al. Haploinsufficiency of GATA-2 perturbs adult hematopoietic stem-cell homeostasis. Blood. 2005;106(2):477–84.

10. Tsai F-Y, Keller G, Kuo FC, Weiss M, Chen J, Rosenblatt M, et al. An early haematopoietic defect in mice lacking the transcription factor GATA-2. Nature. 1994;371(6494):221–6.

11. Rodrigues NP, Boyd AS, Fugazza C, May GE, Guo Y, Tipping AJ, et al. GATA-2 regulates granulocyte-macrophage progenitor cell function. *Blood*, The Journal of the American Society of Hematology. 2008;112(13):4862–73.

12. Li Y, Qi X, Liu B, and Huang H. The STAT5–GATA2 pathway is critical in basophil and mast cell differentiation and maintenance. The Journal of Immunology. 2015;194(9):4328–38.

13. Huang Z, Dore LC, Li Z, Orkin SH, Feng G, Lin S, et al. GATA-2 reinforces megakaryocyte development in the absence of GATA-1. Molecular and cellular biology. 2009;29(18):5168–80.

14. Lulli V, Romania P, Morsilli O, Gabbianelli M, Pagliuca A, Mazzeo S, et al. Overexpression of Ets-1 in human hematopoietic progenitor cells blocks erythroid and promotes megakaryocytic differentiation. Cell Death Differ. 2006;13(7):1064–74.

15. Persons DA, Allay JA, Allay ER, Ashmun RA, Orlic D, Jane SM, et al. Enforced expression of the GATA-2 transcription factor blocks normal hematopoiesis. *Blood*, The Journal of the American Society of Hematology. 1999;93(2):488–99.

16. Tipping AJ, Pina C, Castor A, Hong D, Rodrigues NP, Lazzari L, et al. High GATA-2 expression inhibits human hematopoietic stem and progenitor cell function by effects on cell cycle. *Blood*, The Journal of the American Society of Hematology. 2009;113(12):2661–72.

17. Vicente C, Vazquez I, Conchillo A, Garcia-Sanchez M, Marcotegui N, Fuster O, et al. Overexpression of GATA2 predicts an adverse prognosis for patients with acute myeloid leukemia and it is associated with distinct molecular abnormalities. Leukemia. 2012;26(3):550–4.

18. Menendez-Gonzalez JB, Sinnadurai S, Gibbs A, Thomas L-a, Konstantinou M, Garcia-Valverde A, et al. Inhibition of GATA2 restrains cell proliferation and enhances apoptosis and chemotherapy mediated apoptosis in human GATA2 overexpressing AML cells. Sci Rep. 2019;9(1):12212.

19. Menendez-Gonzalez JB, Strange KE, Bassetto M, Brancale A, Rodrigues NP, and Ferla S. Ligand-based discovery of a novel GATA2 inhibitor targeting acute myeloid leukemia cells. Frontiers in Drug Discovery. 2022;2:1013229.

20. Collin M, Dickinson R, and Bigley V. Haematopoietic and immune defects associated with GATA2 mutation. Br J Haematol. 2015;169(2):173–87.

21. Wlodarski MW, Collin M, and Horwitz MS. Semin Hematol. Elsevier; 2017:81–6.

22. Katsumura KR, Mehta C, Hewitt KJ, Soukup AA, Fraga de Andrade I, Ranheim EA, et al. Human leukemia mutations corrupt but do not abrogate GATA-2 function. Proceedings of the National Academy of Sciences. 2018;115(43):E10109–E18.

23. Hahn CN, Chong C-E, Carmichael CL, Wilkins EJ, Brautigan PJ, Li X-C, et al. Heritable GATA2 mutations associated with familial myelodysplastic syndrome and acute myeloid leukemia. Nat Genet. 2011;43(10):1012–7.

24. Johnson KD, Hsu AP, Ryu M-J, Wang J, Gao X, Boyer ME, et al. Cis-element mutated in GATA2-dependent immunodeficiency governs hematopoiesis and vascular integrity. The Journal of clinical investigation. 2012;122(10):3692–704.

25. Rodrigues NP, Tipping AJ, Wang Z, and Enver T. GATA-2 mediated regulation of normal hematopoietic stem/progenitor cell function, myelodysplasia and myeloid leukemia. The international journal of biochemistry & cell biology. 2012;44(3):457–60.

26. Hsu AP, Sampaio EP, Khan J, Calvo KR, Lemieux JE, Patel SY, et al. Mutations in GATA2 are associated with the autosomal dominant and sporadic monocytopenia and mycobacterial infection (MonoMAC) syndrome. *Blood*, The Journal of the American Society of Hematology. 2011;118(10):2653–5.

27. Dickinson RE, Griffin H, Bigley V, Reynard LN, Hussain R, Haniffa M, et al. Exome sequencing identifies GATA-2 mutation as the cause of dendritic cell, monocyte, B and NK lymphoid deficiency. *Blood*, The Journal of the American Society of Hematology. 2011;118(10):2656–8.

28. Ostergaard P, Simpson MA, Connell FC, Steward CG, Brice G, Woollard WJ, et al. Mutations in GATA2 cause primary lymphedema associated with a predisposition to acute myeloid leukemia (Emberger syndrome). Nat Genet. 2011;43(10):929–31.

29. Bigley V, Haniffa M, Doulatov S, Wang X-N, Dickinson R, McGovern N, et al. The human syndrome of dendritic cell, monocyte, B and NK lymphoid deficiency. J Exp Med. 2011;208(2):227–34.

30. Lim K-C, Hosoya T, Brandt W, Ku C-J, Hosoya-Ohmura S, Camper SA, et al. Conditional Gata2 inactivation results in HSC loss and lymphatic mispatterning. The Journal of clinical investigation. 2012;122(10):3705–17.

31. Kazenwadel J, Betterman KL, Chong C-E, Stokes PH, Lee YK, Secker GA, et al. GATA2 is required for lymphatic vessel valve development and maintenance. The Journal of clinical investigation. 2015;125(8):2979–94.

32. Vinh DC, Patel SY, Uzel G, Anderson VL, Freeman AF, Olivier KN, et al. Autosomal dominant and sporadic monocytopenia with susceptibility to mycobacteria, fungi, papillomaviruses, and myelodysplasia. *Blood*, The Journal of the American Society of Hematology. 2010;115(8):1519–29.

33. West RR, Hsu AP, Holland SM, Cuellar-Rodriguez J, and Hickstein DD. Acquired ASXL1 mutations are common in patients with inherited GATA2 mutations and correlate with myeloid transformation. Haematologica. 2014;99(2):276.

34. Bödör C, Renneville A, Smith M, Charazac A, Iqbal S, Étancelin P, et al. Germ-line GATA2 p. THR354MET mutation in familial myelodysplastic syndrome with acquired monosomy 7 and ASXL1 mutation demonstrating rapid onset and poor survival. Haematologica. 2012;97(6):890.

35. Butko E, Distel M, Pouget C, Weijts B, Kobayashi I, Ng K, et al. Gata2b is a restricted early regulator of hemogenic endothelium in the zebrafish embryo. Development. 2015;142(6):1050–61.

36. Ling K-W, Ottersbach K, Van Hamburg JP, Oziemlak A, Tsai F-Y, Orkin SH, et al. GATA-2 plays two functionally distinct roles during the ontogeny of hematopoietic stem cells. The Journal of experimental medicine. 2004;200(7):871–82.

37. Nováková M, Žaliová M, Suková M, Wlodarski M, Janda A, Froňková E, et al. Loss of B cells and their precursors is the most constant feature of GATA-2 deficiency in childhood myelodysplastic syndrome. Haematologica. 2016;101(6):707.

38. Koegel AK, Hofmann I, Moffitt K, Degar B, Duncan C, and Tubman VN. Acute lymphoblastic leukemia in a patient with MonoMAC syndrome/GATA2 haploinsufficiency. Pediatr Blood Cancer. 2016;63(10):1844–7.

39. Donadieu J, Lamant M, Fieschi C, de Fontbrune FS, Caye A, Ouachee M, et al. Natural history of GATA2 deficiency in a survey of 79 French and Belgian patients. Haematologica. 2018;103(8):1278.

40. Abdelfattah A, Hughes-Davies A, Clayfield L, Menendez-Gonzalez JB, Almotiri A, Alotaibi B, et al. Gata2 haploinsufficiency promotes proliferation and functional decline of hematopoietic stem cells with myeloid bias during aging. Blood Advances. 2021;5(20):4285–90.

41. Abdel-Wahab O, Gao J, Adli M, Dey A, Trimarchi T, Chung YR, et al. Deletion of Asxl1 results in myelodysplasia and severe developmental defects in vivo. J Exp Med. 2013;210(12):2641–59.

42. Charles MA, Saunders TL, Wood WM, Owens K, Parlow A, Camper SA, et al. Pituitary-specific Gata2 knockout: effects on gonadotrope and thyrotrope function. Mol Endocrinol. 2006;20(6):1366–77.

43. Stadtfeld M, and Graf T. Assessing the role of hematopoietic plasticity for endothelial and hepatocyte development by non-invasive lineage tracing. 2005.

44. Adolfsson J, Månsson R, Buza-Vidas N, Hultquist A, Liuba K, Jensen CT, et al. Identification of Flt3+ lympho-myeloid stem cells lacking erythro-megakaryocytic potential: a revised road map for adult blood lineage commitment. Cell. 2005;121(2):295–306.

45. Akashi K, Traver D, Miyamoto T, and Weissman IL. A clonogenic common myeloid progenitor that gives rise to all myeloid lineages. Nature. 2000;404(6774):193–7.

46. Kiel MJ, Yilmaz ÖH, Iwashita T, Yilmaz OH, Terhorst C, and Morrison SJ. SLAM family receptors distinguish hematopoietic stem and progenitor cells and reveal endothelial niches for stem cells. Cell. 2005;121(7):1109–21.

47. Kondo M, Weissman IL, and Akashi K. Identification of clonogenic common lymphoid progenitors in mouse bone marrow. Cell. 1997;91(5):661–72.

48. Oguro H, Ding L, and Morrison SJ. SLAM family markers resolve functionally distinct subpopulations of hematopoietic stem cells and multipotent progenitors. Cell stem cell. 2013;13(1):102–16.

49. Pronk CJ, Rossi DJ, Månsson R, Attema JL, Norddahl GL, Chan CKF, et al. Elucidation of the phenotypic, functional, and molecular topography of a myeloerythroid progenitor cell hierarchy. Cell stem cell. 2007;1(4):428–42.

50. Duran-Struuck R, and Dysko RC. Principles of bone marrow transplantation (BMT): providing optimal veterinary and husbandry care to irradiated mice in BMT studies. Journal of the American Association for Laboratory Animal Science. 2009;48(1):11–22.

51. Kingston RE, Chen CA, and Rose JK. Calcium phosphate transfection. Curr Protoc Mol Biol. 2003;63(1):9.1. -9.1. 11.

52. Schmittgen TD, and Livak KJ. Analyzing real-time PCR data by the comparative CT method. Nat Protoc. 2008;3(6):1101–8.

53. Love MI, Huber W, and Anders S. Moderated estimation of fold change and dispersion for RNA-seq data with DESeq2. Genome Biol. 2014;15:1–21.

54. Benjamini Y, and Hochberg Y. Controlling the false discovery rate: a practical and powerful approach to multiple testing. Journal of the Royal statistical society: series B (Methodological). 1995;57(1):289–300.

55. Subramanian A, Tamayo P, Mootha VK, Mukherjee S, Ebert BL, Gillette MA, et al. Gene set enrichment analysis: a knowledge-based approach for interpreting genome-wide expression profiles. Proceedings of the National Academy of Sciences. 2005;102(43):15545–50.

56. Almotiri A, Alzahrani H, Menendez-Gonzalez JB, Abdelfattah A, Alotaibi B, Saleh L, et al. Zeb1 modulates hematopoietic stem cell fates required for suppressing acute myeloid leukemia. The Journal of clinical investigation. 2021;131(1).

57. Schreck C, Istvánffy R, Ziegenhain C, Sippenauer T, Ruf F, Henkel L, et al. Niche WNT5A regulates the actin cytoskeleton during regeneration of hematopoietic stem cells. J Exp Med. 2017;214(1):165–81.

58. Luis T, Killmann N, and Staal F. Signal transduction pathways regulating hematopoietic stem cell biology: introduction to a series of Spotlight Reviews. Leukemia. 2012;26(1):86–90.

59. de Bruin AM, Demirel Ö, Hooibrink B, Brandts CH, and Nolte MA. Interferon-γ impairs proliferation of hematopoietic stem cells in mice. *Blood*, The Journal of the American Society of Hematology. 2013;121(18):3578–85.

60. Schuettpelz LG, and Link DC. Regulation of hematopoietic stem cell activity by inflammation. Front Immunol. 2013;4:204.

61. Mah L, El-Osta A, and Karagiannis T. γH2AX: a sensitive molecular marker of DNA damage and repair. Leukemia. 2010;24(4):679–86.

62. Yang J, Zhang L, Yu C, Yang X-F, and Wang H. Monocyte and macrophage differentiation: circulation inflammatory monocyte as biomarker for inflammatory diseases. Biomarker research. 2014;2:1–9.

63. Takai J, Shimada T, Nakamura T, Engel JD, and Moriguchi T. Gata2 heterozygous mutant mice exhibit reduced inflammatory responses and impaired bacterial clearance. IScience. 2021;24(8).

64. Wang J, Li Z, He Y, Pan F, Chen S, Rhodes S, et al. Loss of Asxl1 leads to myelodysplastic syndrome–like disease in mice. Blood, The Journal of the American Society of Hematology. 2014;123(4):541–53.

65. Fujino T, Goyama S, Sugiura Y, Inoue D, Asada S, Yamasaki S, et al. Mutant ASXL1 induces age-related expansion of phenotypic hematopoietic stem cells through activation of Akt/mTOR pathway. Nature communications. 2021;12(1):1826.

66. Lawson H, van de Lagemaat LN, Barile M, Tavosanis A, Durko J, Villacreces A, et al. CITED2 coordinates key hematopoietic regulatory pathways to maintain the HSC pool in both steady-state hematopoiesis and transplantation. Stem cell reports. 2021;16(11):2784–97.

67. Micol J-B, Pastore A, Inoue D, Duployez N, Kim E, Lee SC-W, et al. ASXL2 is essential for haematopoiesis and acts as a haploinsufficient tumour suppressor in leukemia. Nature communications. 2017;8(1):15429.

68. Zhang P, Chen Z, Li R, Guo Y, Shi H, Bai J, et al. Loss of ASXL1 in the bone marrow niche dysregulates hematopoietic stem and progenitor cell fates. Cell discovery. 2018;4(1):4.

69. Chigaev A. Does aberrant membrane transport contribute to poor outcome in adult acute myeloid leukemia? Front Pharmacol. 2015;6:134.

70. Raaijmakers M. ATP-binding-cassette transporters in hematopoietic stem cells and their utility as therapeutical targets in acute and chronic myeloid leukemia. Leukemia. 2007;21(10):2094–102.

71. Hwang CY, Choe W, Yoon K-S, Ha J, Kim SS, Yeo E-J, et al. Molecular mechanisms for ketone body metabolism, signaling functions, and therapeutic potential in cancer. Nutrients. 2022;14(22):4932.

72. Chiarella E, Nisticò C, Di Vito A, Morrone HL, and Mesuraca M. Targeting of mevalonate-isoprenoid pathway in acute myeloid leukemia cells by bisphosphonate drugs. Biomedicines. 2022;10(5):1146.

73. Pei S, Minhajuddin M, Callahan KP, Balys M, Ashton JM, Neering SJ, et al. Targeting aberrant glutathione metabolism to eradicate human acute myelogenous leukemia cells. J Biol Chem. 2013;288(47):33542–58.

74. Ohanian M, Rozovski U, Kanagal-Shamanna R, Abruzzo LV, Loghavi S, Kadia T, et al. MYC protein expression is an important prognostic factor in acute myeloid leukemia. Leuk Lymphoma. 2019;60(1):37–48.

75. Ren B, Cam H, Takahashi Y, Volkert T, Terragni J, Young RA, et al. E2F integrates cell cycle progression with DNA repair, replication, and G2/M checkpoints. Genes Dev. 2002;16(2):245–56.

76. Sabbah R, Saadi S, Shahar-Gabay T, Gerassy S, Yehudai-Resheff S, and Zuckerman T. Abnormal adipogenic signaling in the bone marrow mesenchymal stem cells contributes to supportive microenvironment for leukemia development. Cell Communication and Signaling. 2023;21(1):277.

77. Ochi Y, and Ogawa S. Chromatin-spliceosome mutations in acute myeloid leukemia. Cancers (Basel). 2021;13(6):1232.

78. Féral K, Jaud M, Philippe C, Di Bella D, Pyronnet S, Rouault-Pierre K, et al. ER stress and unfolded protein response in leukemia: friend, foe, or both? Biomolecules. 2021;11(2):199.

79. Brendolan A, and Russo V. Targeting cholesterol homeostasis in hematopoietic malignancies. *Blood*, The Journal of the American Society of Hematology. 2022;139(2):165–76.

80. Soltani M, Zhao Y, Xia Z, Ganjalikhani Hakemi M, and Bazhin AV. The importance of cellular metabolic pathways in pathogenesis and selective treatments of hematological malignancies. Front Oncol. 2021;11:767026.

81. Jones CL, Stevens BM, D’Alessandro A, Culp-Hill R, Reisz JA, Pei S, et al. Cysteine depletion targets leukemia stem cells through inhibition of electron transport complex II. *Blood*, The Journal of the American Society of Hematology. 2019;134(4):389–94.

82. Cunningham A, Erdem A, Alshamleh I, Geugien M, Pruis M, Pereira-Martins DA, et al. Dietary methionine starvation impairs acute myeloid leukemia progression. *Blood*, The Journal of the American Society of Hematology. 2022;140(19):2037–52.

83. Mishra SK, Millman SE, and Zhang L. Metabolism in acute myeloid leukemia: mechanistic insights and therapeutic targets. Blood. 2023;141(10):1119–35.

84. Matsumoto N, Yoneda-Kato N, Iguchi T, Kishimoto Y, Kyo T, Sawada H, et al. Elevated MLF1 expression correlates with malignant progression from myelodysplastic syndrome. Leukemia. 2000;14(10):1757–65.

85. Rochette L, Zeller M, Cottin Y, and Vergely C. Insights into mechanisms of GDF15 and receptor GFRAL: therapeutic targets. Trends Endocrinol Metab. 2020;31(12):939–51.

86. Hu F, Chen Sl, Dai Yj, Wang Y, Qin Zy, Li H, et al. Identification of a metabolic gene panel to predict the prognosis of myelodysplastic syndrome. Journal of Cellular and Molecular Medicine. 2020;24(11):6373–84.

87. Liu J, Xiao Q, Xiao J, Niu C, Li Y, Zhang X, et al. Wnt/β-catenin signalling: function, biological mechanisms, and therapeutic opportunities. Signal transduction and targeted therapy. 2022;7(1):3.

88. Pellagatti A, Armstrong RN, Steeples V, Sharma E, Repapi E, Singh S, et al. Impact of spliceosome mutations on RNA splicing in myelodysplasia: dysregulated genes/pathways and clinical associations. *Blood*, The Journal of the American Society of Hematology. 2018;132(12):1225–40.

89. Soukup AA, Zheng Y, Mehta C, Wu J, Liu P, Cao M, et al. Single-nucleotide human disease mutation inactivates a blood-regenerative GATA2 enhancer. The Journal of clinical investigation. 2019;129(3):1180–92.

90. Suzuki M, Kobayashi-Osaki M, Tsutsumi S, Pan X, Ohmori Sy, Takai J, et al. GATA factor switching from GATA 2 to GATA 1 contributes to erythroid differentiation. Genes Cells. 2013;18(11):921–33.

91. Guo G, Luc S, Marco E, Lin T-W, Peng C, Kerenyi MA, et al. Mapping cellular hierarchy by single-cell analysis of the cell surface repertoire. Cell stem cell. 2013;13(4):492–505.

92. Manso BA, Rodriguez y Baena A, and Forsberg EC. From Hematopoietic Stem Cells to Platelets: Unifying Differentiation Pathways Identified by Lineage Tracing Mouse Models. Cells. 2024;13(8):704.

93. Bruns I, Lucas D, Pinho S, Ahmed J, Lambert MP, Kunisaki Y, et al. Megakaryocytes regulate hematopoietic stem cell quiescence through CXCL4 secretion. Nat Med. 2014;20(11):1315–20.

94. Nakamura-Ishizu A, Takubo K, Fujioka M, and Suda T. Megakaryocytes are essential for HSC quiescence through the production of thrombopoietin. Biochemical and biophysical research communications. 2014;454(2):353–7.

95. Zhao M, Perry JM, Marshall H, Venkatraman A, Qian P, He XC, et al. Megakaryocytes maintain homeostatic quiescence and promote post-injury regeneration of hematopoietic stem cells. Nat Med. 2014;20(11):1321–6.

96. Wu Z, Gao S, Diamond C, Kajigaya S, Chen J, Shi R, et al. Sequencing of RNA in single cells reveals a distinct transcriptome signature of hematopoiesis in GATA2 deficiency. Blood Advances. 2020;4(12):2702–16.

97. Calvo KR, Vinh DC, Maric I, Wang W, Noel P, Stetler-Stevenson M, et al. Myelodysplasia in autosomal dominant and sporadic monocytopenia immunodeficiency syndrome: diagnostic features and clinical implications. Haematologica. 2011;96(8):1221.

98. Biechonski S, Yassin M, and Milyavsky M. DNA-damage response in hematopoietic stem cells: an evolutionary trade-off between blood regeneration and leukemia suppression. Carcinogenesis. 2017;38(4):367–77.

99. Moehrle BM, and Geiger H. Aging of hematopoietic stem cells: DNA damage and mutations? Exp Hematol. 2016;44(10):895–901.

100. Niedernhofer LJ. DNA repair is crucial for maintaining hematopoietic stem cell function. DNA Repair. 2008;7(3):523–9.

101. Mohrin M, Bourke E, Alexander D, Warr MR, Barry-Holson K, Le Beau MM, et al. Hematopoietic stem cell quiescence promotes error-prone DNA repair and mutagenesis. Cell stem cell. 2010;7(2):174–85.

102. Zhou T, Chen P, Gu J, Bishop AJ, Scott LM, Hasty P, et al. Potential relationship between inadequate response to DNA damage and development of myelodysplastic syndrome. Int J Mol Sci. 2015;16(1):966–89.

103. Pietras EM, Warr MR, and Passegué E. Cell cycle regulation in hematopoietic stem cells. J Cell Biol. 2011;195(5):709–20.

104. Avagyan S, Henninger J, Mannherz W, Mistry M, Yoon J, Yang S, et al. Resistance to inflammation underlies enhanced fitness in clonal hematopoiesis. Science. 2021;374(6568):768–72.

105. Mannava S, Grachtchouk V, Wheeler LJ, Im M, Zhuang D, Slavina EG, et al. Direct role of nucleotide metabolism in C-MYC-dependent proliferation of melanoma cells. Cell cycle. 2008;7(15):2392–400.

106. Dang CV, O’Donnell KA, Zeller KI, Nguyen T, Osthus RC, and Li F. Semin Cancer Biol. Elsevier; 2006:253–64.

