## Supplementary Figures 1-7 for "Impaired DNA damage response and inflammatory signalling underpins hematopoietic stem cell defects in *Gata2* haploinsufficiency"

### Slide 1
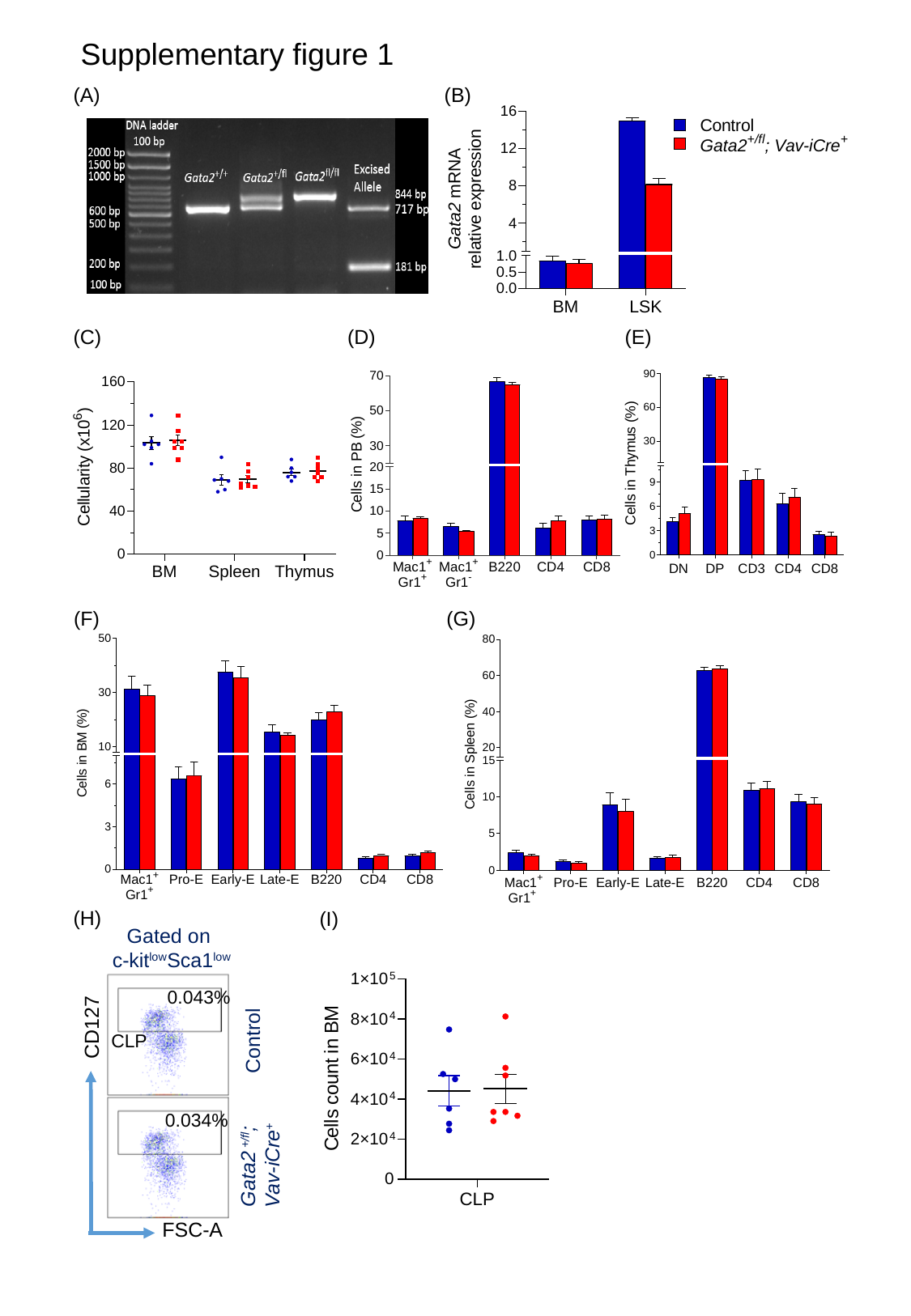

Supplementary figure 1
(A)
(B)
(D)
(C)
(E)
(G)
(F)
(H)
(I)
Gated on
c-kit^low^Sca1^low^
0.043%
CD127
Control
CLP
0.034%
Gata2 ^+/fl^;
Vav-iCre^+^
FSC-A

### Slide 2
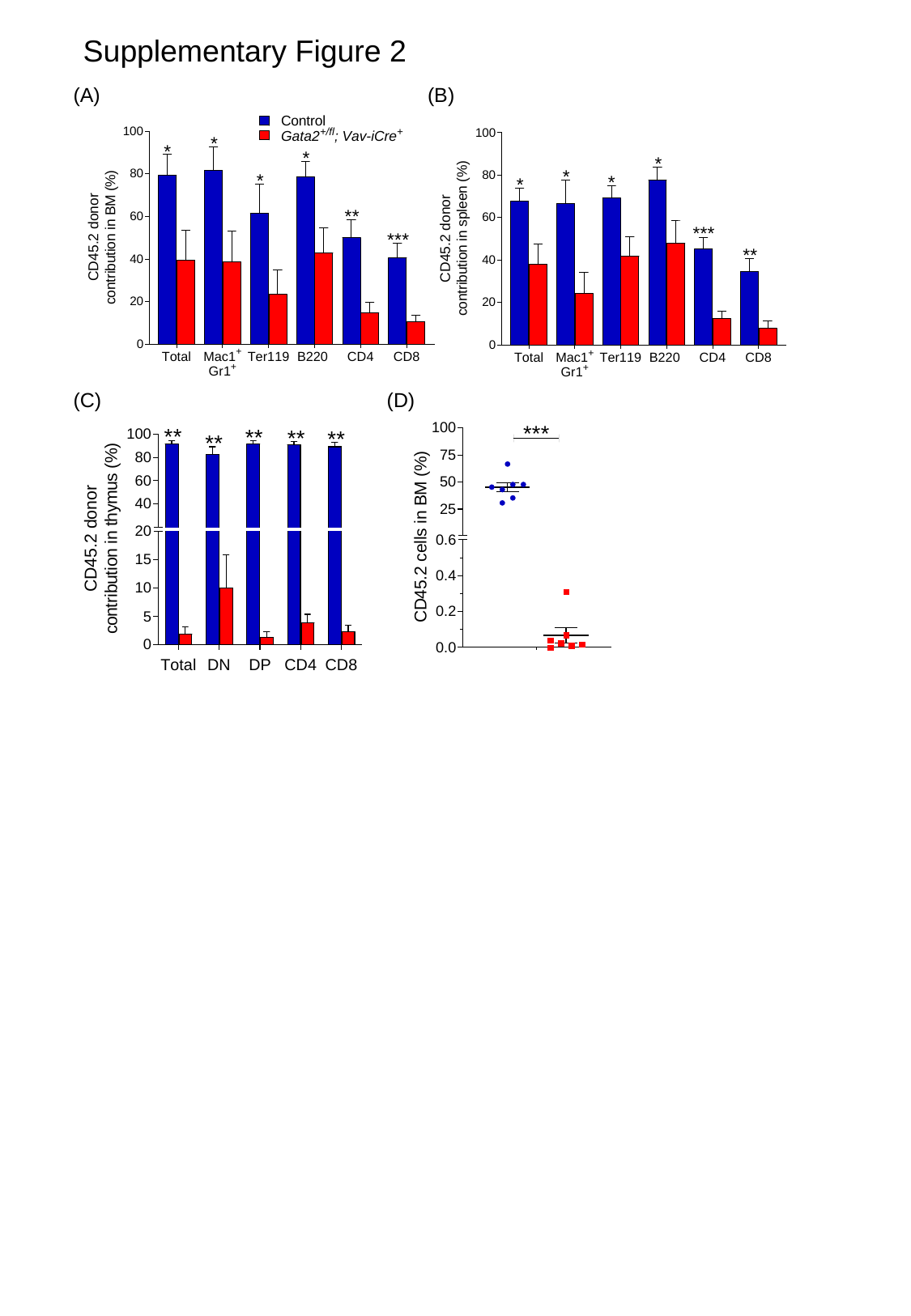

Supplementary Figure 2
(A)
(B)
(C)
(D)

### Slide 3
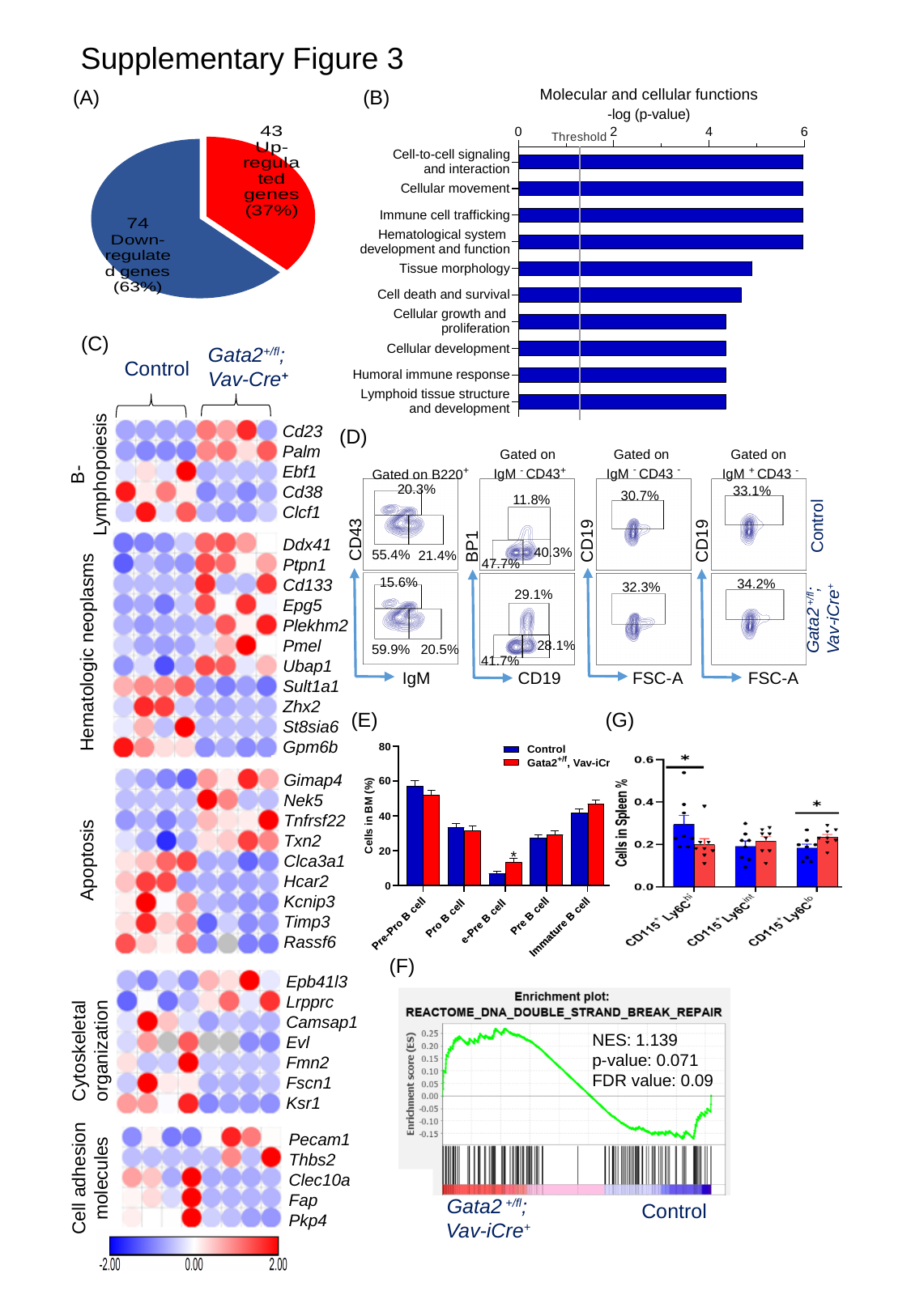

Supplementary Figure 3
(B)
(A)
#### Chart
| Category | Sales |
|---|---|
| Up-regulated genes | 43.0 |
| Down-regulated genes | 74.0 |(C)
Gata2^+/fl^;
Vav-Cre^+^
Control
Cd23
Palm
Ebf1
Cd38
Clcf1
(D)
Gated on
IgM ^-^ CD43 ^-^
Gated on
 IgM ^-^ CD43^+^
Gated on
 IgM ^+^ CD43 ^-^
B-
Lymphopoiesis
Gated on B220^+^
20.3%
33.1%
30.7%
11.8%
Control
CD43
CD19
BP1
CD19
Ddx41
Ptpn1
Cd133
Epg5
Plekhm2
Pmel
Ubap1
Sult1a1
Zhx2
St8sia6
Gpm6b
40.3%
55.4%
21.4%
47.7%
15.6%
34.2%
32.3%
29.1%
Gata2 ^+/fl^;
Vav-iCre^+^
28.1%
59.9%
Hematologic neoplasms
20.5%
41.7%
 IgM
FSC-A
 CD19
FSC-A
(E)
(G)
Gimap4
Nek5
Tnfrsf22
Txn2
Clca3a1
Hcar2
Kcnip3
Timp3
Rassf6
Apoptosis
(F)
Epb41l3
Lrpprc
Camsap1
Evl
Fmn2
Fscn1
Ksr1
Cytoskeletal
organization
NES: 1.139
p-value: 0.071
FDR value: 0.09
Pecam1
Thbs2
Clec10a
Fap
Pkp4
Cell adhesion
 molecules
Gata2 ^+/fl^;
Vav-iCre^+^
Control

### Slide 4
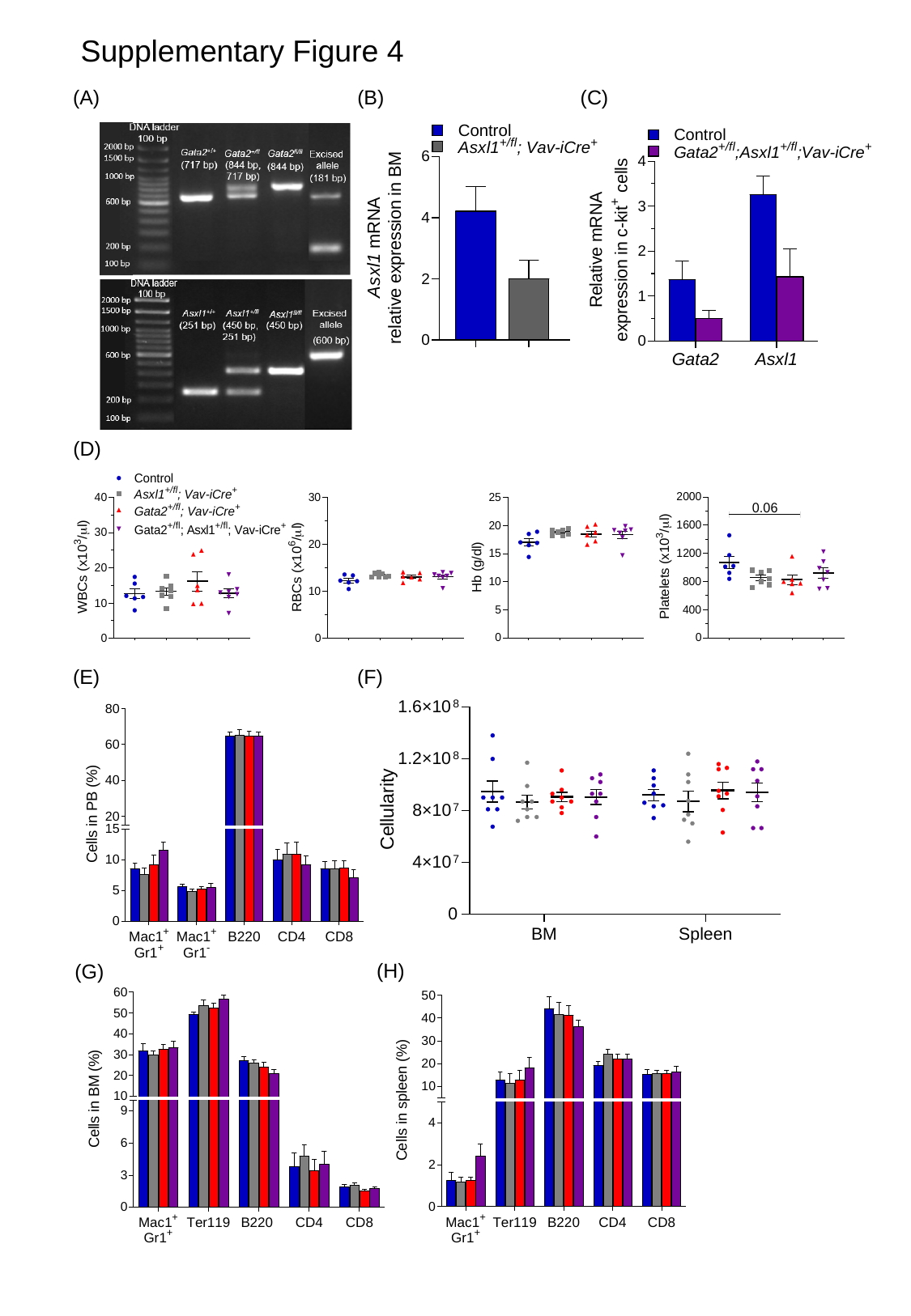

Supplementary Figure 4
(A)
(B)
(C)
(D)
(E)
(F)
(H)
(G)

### Slide 5
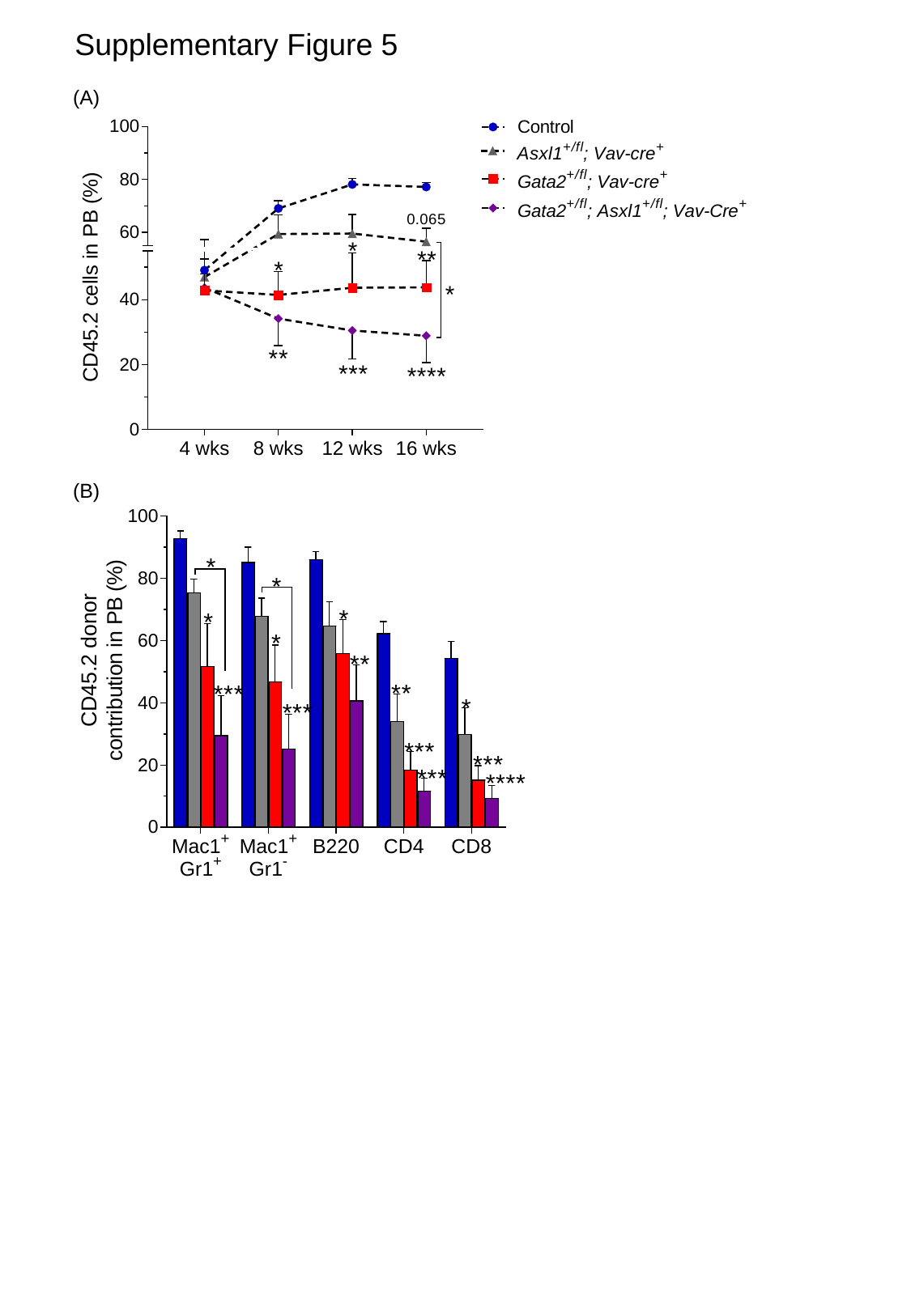

Supplementary Figure 5
(A)
(B)

### Slide 6
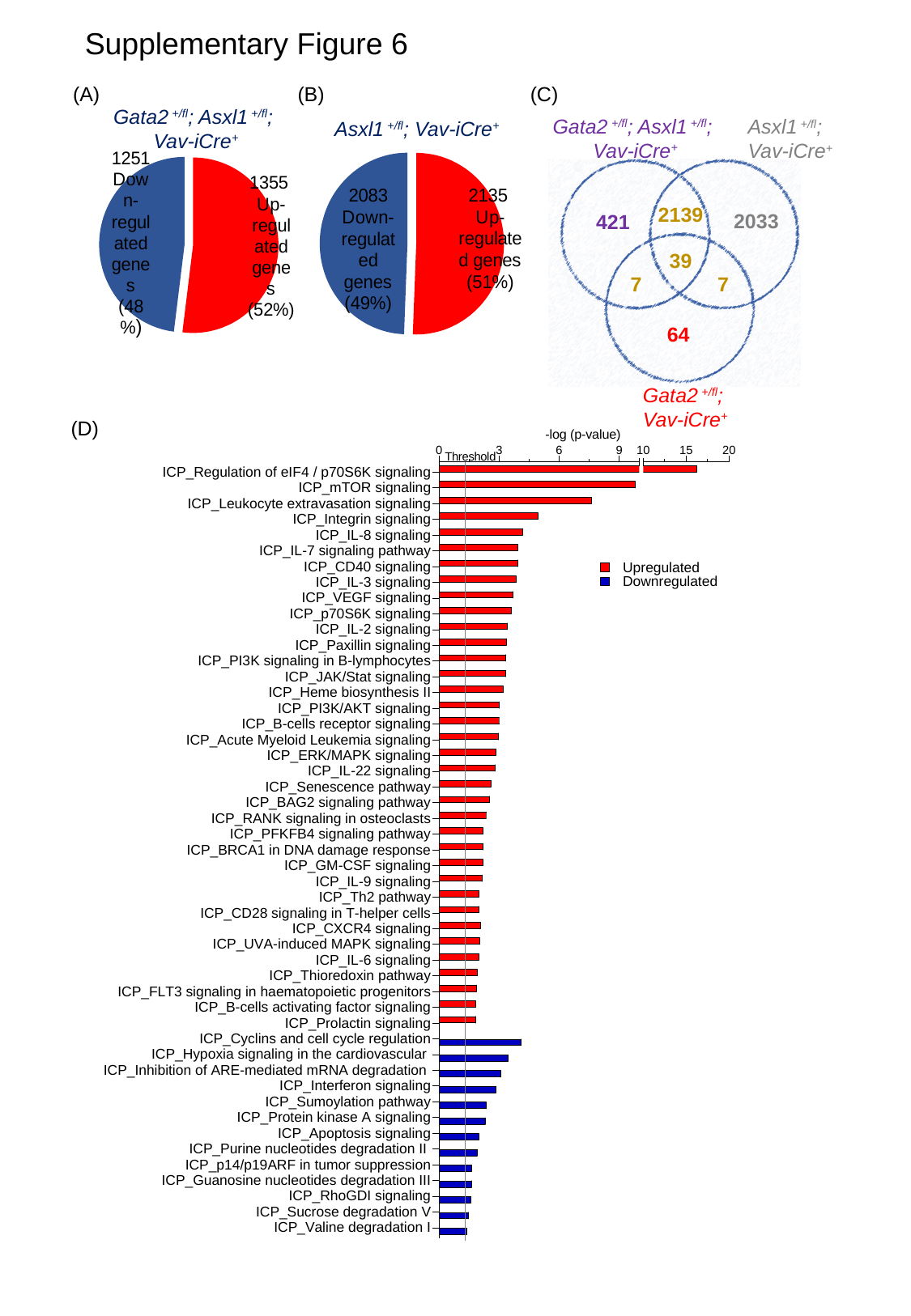

Supplementary Figure 6
(A)
(B)
(C)
Gata2 ^+/fl^; Asxl1 ^+/fl^;
 Vav-iCre^+^
Gata2 ^+/fl^; Asxl1 ^+/fl^;
 Vav-iCre^+^
Asxl1 ^+/fl^;
Vav-iCre^+^
Asxl1 ^+/fl^; Vav-iCre^+^
#### Chart
| Category | Significant genes |
|---|---|
| Up-regulated genes | 1355.0 |
| Down-regulated genes | 1251.0 |
#### Chart
| Category | Significant genes |
|---|---|
| Up-regulated genes | 2135.0 |
| Down-regulated genes | 2083.0 |
2139
2033
421
39
7
7
64
Gata2 ^+/fl^;
Vav-iCre^+^
(D)

### Slide 7
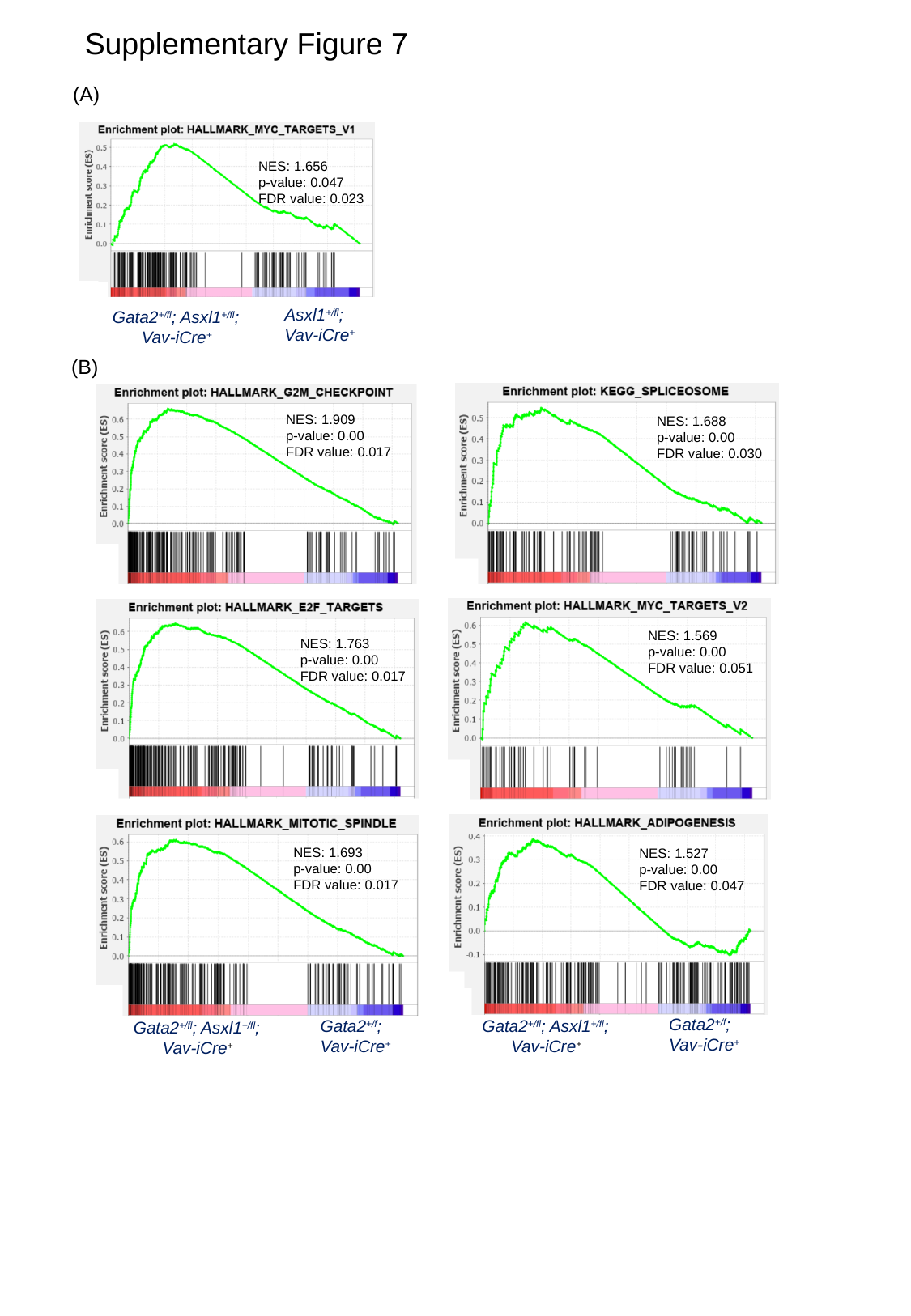

Supplementary Figure 7
(A)
NES: 1.656
p-value: 0.047
FDR value: 0.023
Asxl1^+/fl^;
Vav-iCre^+^
Gata2^+/fl^; Asxl1^+/fl^;
Vav-iCre^+^
(B)
NES: 1.909
p-value: 0.00
FDR value: 0.017
NES: 1.688
p-value: 0.00
FDR value: 0.030
NES: 1.569
p-value: 0.00
FDR value: 0.051
NES: 1.763
p-value: 0.00
FDR value: 0.017
NES: 1.693
p-value: 0.00
FDR value: 0.017
NES: 1.527
p-value: 0.00
FDR value: 0.047
Gata2^+/f^;
Vav-iCre^+^
Gata2^+/f^;
Vav-iCre^+^
Gata2^+/fl^; Asxl1^+/fl^;
Vav-iCre^+^
Gata2^+/fl^; Asxl1^+/fl^;
Vav-iCre^+^
